# Direct Reprogramming of Non-limb Fibroblasts to Cells with Properties of Limb Progenitors

**DOI:** 10.1101/2021.10.01.462632

**Authors:** Yuji Atsuta, Changhee Lee, Alan R. Rodrigues, Charlotte Colle, Reiko R. Tomizawa, Ernesto G. Lujan, Patrick Tschopp, Joshua M. Gorham, Jean-Pierre Vannier, Christine E. Seidman, Jonathan G. Seidman, Olivier Pourquié, Clifford J. Tabin

## Abstract

The early limb bud consists of mesenchymal progenitors (limb progenitors) derived from the lateral plate mesoderm (LPM) that produce most of the tissues of the mature limb bud. The LPM also gives rise to the mesodermal components of the trunk, flank and neck. However, the mesenchymal cells generated at these other axial levels cannot produce the variety of cell types found in the limb bud, nor can they be directed to form a patterned appendage-like structure, even when placed in the context of the signals responsible for organizing the limb bud. Here, by taking advantage of a direct reprogramming approach, we find a set of factors (Prdm16, Zbtb16, and Lin28) normally expressed in the early limb bud, that are capable of imparting limb progenitor-like properties to non-limb fibroblasts. Cells reprogrammed by these factors show similar gene expression profiles, and can differentiate into similar cell types, as endogenous limb progenitors. The further addition of Lin41 potentiates proliferation of the reprogrammed cells while suppressing differentiation. These results suggest that these same four key factors may play pivotal roles in the specification of endogenous limb progenitors.

## INTRODUCTION

Limb bud progenitors originate from the somatopleural layer of the LPM, a continuous epithelium lining the embryonic coelom. Limb progenitors emerge through localized epithelial-to-mesenchymal transition (EMT) at limb forming levels (Gros and Tabin, 2014). Limb progenitors will ultimately give rise to the majority of tissues present in the mature patterned limb including cartilage, bone, tendon, ligament, muscle connective tissue and dermis; whereas somatopleural LPM at other axial levels, such as neck and flank mesenchyme, will only form dermis. Moreover, limb progenitors are organized within the limb bud in response to limb-patterning morphogenic signals, while LPM-derived cells from other axial levels are refractory to them (Takeuchi et al., 2003). It has, however, remained unclear what gene, or genes, are responsible for specifying limb progenitors and imparting them with limb-specific traits.

In previous studies, direct cellular reprogramming has been used to induce a variety of tissue progenitor populations, such as neural progenitors, cardiomyocytes and hepatocytes, from terminally differentiated fibroblasts (Vierbuchen et al., 2010). These studies not only set the stage for future therapeutic applications, but they have also proven important, in and of themselves, for identifying developmental regulators of embryonic progenitor states (Takahashi and Yamanaka, 2015). For example, the reprogramming factors first shown to be capable of inducing pluripotent stem cells (Oct3/4, Sox2, Klf4 and c-Myc) were subsequently shown to regulate the endogenous developmental signaling network defining mouse embryonic stem cells (Lin et al., 2008).

To understand what it really means to be a limb progenitor, we set out to identify a set of factors expressed ubiquitously in the early limb field, that might be capable of establishing and maintaining the unique transcriptional characteristics and differentiation potential of limb progenitors. To that end, we took a reprogramming approach, reasoning that a full set of the factors giving early limb progenitors their unique properties might be sufficient to convert non-limb mouse embryonic fibroblasts into cells with properties of limb progenitors.

We started with 18 candidate factors expressed in early limb progenitors. We overexpressed these factors via viral vectors in three-dimensional (3D) culture conditions optimized for maintaining legitimate limb progenitors. This pool of 18 factors was, indeed, able to robustly induce expression of limb progenitor marker genes in mouse embryonic non-limb fibroblasts. Winnowing the candidates responsible for this activity, we ultimately found that, a combination of two transcription factors, Prdm16 and Zbtb16, plus an RNA-binding protein, Lin28a, suffice to reprogram non-limb fibroblasts into a limb progenitor-like state (reprogrammed limb progenitor-like cells, hereafter rLPCs). Moreover, the further addition of Lin41 (also known as Trim71), boosts proliferation of rLPCs, by antagonizing translation of Egr1, a pro-differentiation factor for limb progenitors. The limb progenitor-like state of the rLPCs was validated at a transcriptional level, and through in vitro and in vivo differentiation assays. While our initial analysis was carried out with murine cells, we further show that adult human fibroblasts can similarly be converted to rLPCs with the same set of factors used for mouse cell reprogramming, suggestive of conservation of the genetic program for limb bud initiation across vertebrates. Taken together, the reprogramming factors identified here are capable of conferring non-limb cells with limb progenitor specific traits, suggesting that these factors might similarly initiate developmental networks that define the endogenous early limb progenitors as they emerge from the LPM.

## Results

### Optimization of culture conditions for early mouse limb bud progenitors

Prior to embarking on a reprogramming strategy, we needed to establish culture conditions capable of sustaining authentic limb progenitors, to assure that putative reprogrammed limb progenitor-like cells would be able to expand into colonies while maintaining a limb progenitor-like identity. 3D-culture systems mimicking physiological conditions have been used to support expansion of primary progenitor populations such as neural and nephron progenitor cells (Madl et al., 2017; Li et al., 2016), as well as for cellular reprogramming of iPSCs (Caiazzo, 2016). To mimic the early limb bud extracellular environment, we exploited hydrogel scaffolds made from high molecular weight hyaluronic acid (HA) and adipic acid dihydrazide crosslinkers. HA is a large glycosaminoglycan that is known to be a major component of the extracellular matrix (ECM) of the developing limb buds (Li et al., 2007).

In a previous study, we showed that treating cultured chick limb bud cells with a combination of Wnt3a, Fgf8 and retinoic acid (RA) maintained them in a progenitor state for 48 hours (Cooper et al., 2011). Here, we utilized CHIR99021, a GSK3 inhibitor in place of Wnt3a. We compared the effect of these factors on mouse limb progenitors, cultured within a 3D-HA-gel scaffold with those maintained in two-dimensional culture on polystyrene plastic. To provide a readout for maintenance of a limb progenitor identity, we harvested limb bud progenitors from E9.5 *Prx1*-GFP reporter mice (*Prx1*-CreER-ires-GFP)(Kawanami et al., 2009), in which GFP activity is specifically seen in the limb buds (Fig. 1A). While the GFP signal was maintained in 2-D culture conditions for the first 48 hours, there was no stimulation of cell proliferation (Fig S1A), and the GFP activity was rapidly lost at later time points. In contrast, under 3-D HA-gel conditions, the plated cells expanded over 20-fold during this time (Fig. 1A, B). With subsequent culture, however, the cell number diminished. Moreover, the expression of three different early limb bud markers, *Prx1*-GFP, Lhx2 and Sall4, were only maintained for the first 2 days of culture (Fig. 1C and S1B) (Rodriguez-Esteban et al., 1998). Reasoning that the loss of limb progenitors could be due to differentiation, cell death, or both, we added Y-27632, a Rho-associated kinase inhibitor (as this factor is known to suppress dissociation-associated cell death of stem/progenitor cells) (Watanabe et al., 2007); and SB431542, a TGFβ/BMP antagonist (as TGFβ and BMP act as pro-differentiation factors for tendons and cartilage, respectively) (Healy et al., 1999). Media supplemented with this combination of CHIR, Fgf8, RA, SB431542 and Y-27632 greatly increased proliferation of limb progenitors (Fig. 1B). Moreover, 49.2% of the cultured cells in the HA-gels remained *Prx*GFP^+^/Lhx2^+^/Sall4^+^-positive for at least 8 days (Fig. 1A, C and S1B). To see if this set of factors could maintain the differentiation potential of cultured limb progenitors, GFP-expressing chicken limb progenitors were kept in a 3D-HA gel supplemented by these factors for 8 days, and were then grafted into host limb buds (Chapman et al., 2005). When observed 5 days later, the transplanted GFP-chick cells were integrated into both cartilage expressing Sox9^+^ (Fig. S2A), and muscle associated tendon, expressing Collagen I (Fig. S2B) indicating that limb progenitors cultured in the 3D-HA-gel in the presence of the defined set of factors maintained their potency to differentiate into limb tissue types.

**Figure 1.**
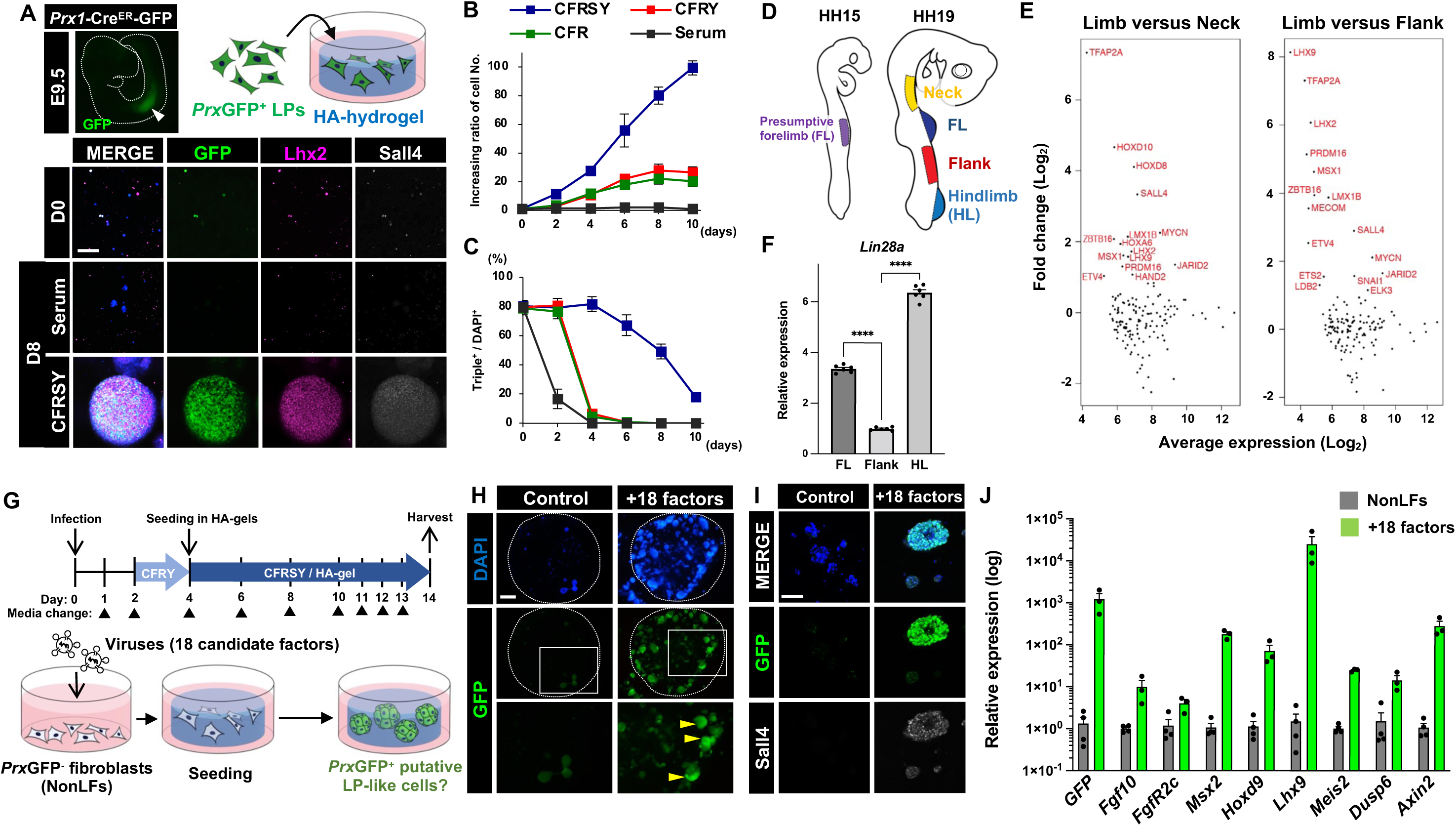
Overexpression of the factors that are present specifically in the limb bud induces expression of limb progenitor genes in non-limb fibroblasts. (A) Optimization of culture conditions for forelimb (FL) progenitors from *Prx1*-GFP mouse embryos (*Prx*GFP^+^ LPs) by using hyaluronan (HA)-based hydrogels. The cultured LPs were stained with antibodies for GFP (green), Lhx2 (magenta) and Sall4 (white). Serum media was DMEM containing 10% FBS, and CFRSY media contained Chir99021 (3 μM), Fgf8 (150 ng/ml), Retinoic acid (25 nM), SB431542 (5 μM) and Y-27632 (10 μM). (B) Increasing ratio of cell number. Cell numbers in Day0 samples of each condition were counted immediately after seeding, and were considered as ratio 1. (C) Percentages for *Prx*GFP/Lhx2/Sall4-triple positive cells in cultures. (D) Schematics of HH stage 15 and HH19 chicken embryos. Regions of embryos that were used for transcriptomic analyses are labeled. (E) Differential expression analyses (MA-plot) of core gene set. Limb expression (average of FL and hindlimb [HL]) over neck or flank expression. Labeled points indicate genes with greater than two-fold overexpression in limb tissue. (F) *Lin28a* mRNA expression levels in FL, flank and HL of E9.5 mouse embryos were measured by qPCR (n = 6 for each). (G) Diagrams illustrating procedures of the reprogramming experiment. Retrovirus particles carrying each factor of 18 candidates were pooled and used to infect non-limb *Prx*GFP-negative fibroblasts (NonLFs) at Day0. After infection, the media was replaced with CFRY (Day2-4), subsequently with CFRSY (Day4-14). The infected NonLFs were seeded in HA-gels at Day4. (H) The cells infected with no virus or 18 viruses carrying candidate factors were visualized by DAPI (blue). Dashed lines indicate outer edge of the hydrogel. Induced *Prx*GFP signals were seen in cell clusters (yellow arrowheads). (I) Magnified images of the cell clusters. Sall4 proteins were observed in *Prx*GFP positive cells. (J) Relative expression levels of *GFP*, *Fgf10*, *FgfR2c*, *Msx2*, *Hoxd9*, *Lhx9*, *Meis2*, *Dusp6* and *Axin2* were quantified by qPCR (n = 4 for NonLFs, n = 3 for +18 factors). \*\*\*\**p* < 0.0001, one-way ANOVA. Error bars represent SD. Scale bars, 100 μm in (A) and (I), 1 mm in (H).

Finally, we asked whether other culture matrices besides HA could maintain limb progenitors in the presence of CHIR99021, Fgf8, RA, SB431542 and Y-27632. Several scaffolds we tested failed to do so, however we discovered that limb bud cells plated onto Matrigel grew to a similar extent, and maintained expression of limb-specific markers, equivalent to those seeded into the HA scaffold (Fig. S3A-C). Accordingly, the HA and Matrigel systems were used interchangeably in subsequent experiments as noted below (being careful to always compare to controls cultured in the same matrix).

### Identification of candidate genes for specification of limb progenitor identity

To generate a list of candidate transcription factors potentially involved in early limb fate specification, we used RNA-seq to identify genes expressed exclusively in the early chick limb fields. We harvested the forelimb and hindlimb buds of HH17-19 embryos, as well as presumptive neck and flank mesenchyme from HH19-20 embryos (Fig. 1D). Additionally, we profiled the epithelial lateral plate mesoderm prior to forelimb bud emergence (HH15; Fig. 1D). The transcriptional profiles of these tissues were compared in a principal component analysis (PCA). The first and second PC accounted for 48% and 28% of the variance in the five data sets. When plotted in the principal component space, the forelimb and hindlimb bud tissues clustered together tightly (Fig. S4A). PC1 separates the remaining three tissues from the limb tissues while PC2 separates epithelial lateral plate and neck mesenchyme from the limb tissues (Fig. S4A). To determine the key drivers of this separation in PC space, the top 100 genes contributing to each principal component were used in a gene set enrichment analysis. For both PC1 and PC2, the top five most significant classes of gene function were related to transcriptional regulation (Fig. S4B), suggesting that the drivers of difference between limb and non-limb lateral plate mesenchyme are transcription factors. We then intersected our existing mouse hindlimb bud transcriptional data set (Tschopp et al., 2014) with our chick data to generate an evolutionarily conserved set of candidate genes we could use in a reprogramming assay. Of the 1806 transcriptional regulators in the mouse genome, 303 are expressed at appreciable levels in the mouse hindlimb. Of these 303 genes, 142 are co-expressed in both the chick forelimb and hindlimb. Of this core set of 142 transcription factors, co-factors and chromatin remodelers, we particularly were interested in those that were differentially expressed relative to the neck and/or flank mesenchyme. Only 15 of the 142 factors were more than two-fold over-expressed in the limb as compared to the neck and 16 were more than two-fold overexpressed when compared to the flank (Fig. 1E). Among those genes, we excluded Lhx9 and Hoxa6 as these genes were deemed potentially redundant to Lhx2 and other Hox genes, respectively. Sall4 was replaced with Sall1, a multi-zinc finger transcription factor that functions redundantly with Sall4 (Bohm et al., 2008), because a reliable antibody against Sall4 was available, which could be considered as a proxy for the reprogramming. Lmx1b was withdrawn because it specifies only the dorsal compartment of the limb field (Chen et al., 1998), and Snai1/2 was removed from the list because limb-specific double mutants show no defect in limb bud formation (Chen and Gridley, 2013). In addition, we included several genes such as Tbx5 and Pbx2, which were not differentially expressed relative to the flank tissue, but were expressed in both the chicken and mouse limb progenitors, and had been previously implicated functionally as being important for limb bud outgrowth (Takeuchi et al., 2003; Capellini et al., 2006).

Finally, we added Lin28a to the list. Lin28a is a highly conserved RNA-binding protein, the major function of which is to bind nascent *let-7* micro RNA in order to block its biogenesis (Viswanathan et al., 2008). Lin28a plays roles in regulating development and pluripotency (Tsialikas and Romer-Seibert, 2015), and is known as one of the iPSC reprogramming factors (Yu et al., 2007). Of note, expression of *Lin28a* mRNA has been specifically seen in early limb buds, in both mouse and chicken embryos, and its expression is downregulated as limb development progresses (Buganim et al., 2014). Moreover, we observe a relatively higher expression level of Lin28a in mouse limb buds than in the flank lateral plate mesoderm (Fig. 1F). Taken together, this generated a list of 18 candidate reprogramming factors.

### Overexpression of candidate genes specifically expressed in early limb buds activates expression of limb progenitor genes in non-limb fibroblasts

We isolated GFP-negative fibroblasts from the non-limb regions of E13.5 *Prx1*-GFP transgenic embryos. These non-limb fibroblasts were infected with pooled retroviruses transducing our 18 candidate factors, and were cultured under the conditions optimized for legitimate limb progenitors (Fig. 1G). Taking advantage of the limb-specific GFP activity as an indicator of reprogramming, we asked if the pooled 18 candidate factors could induce GFP expression in non-limb fibroblasts. Indeed, 14 days after infection, the emergence of GFP positive cells became apparent. Of interest, a fraction of the GFP positive cells formed clusters reminiscent of freshly harvested limb progenitors cultured in the same conditions (Fig. 1A, H). While the *Prx1* promoter strongly drives expression in limb buds, it is also expressed in some other regions of the embryo, such as the head mesoderm. Thus, we examined expression of other limb progenitor marker genes as well (Fig. 1I, J). We observed induction of increased Sall4 protein levels by immunohistochemistry (Fig. 1I), as well as increased transcript levels of other limb progenitor markers (*Prx1-GFP*, *Fgf10*, *FgfR2c*, *Msx2*, *Hoxd9*, *Lhx9*, *Meis2*, *Dusp6* and *Axin2*) measured by qPCR (Fig. 1J). Strikingly, each of these markers was upregulated in infected cells relative to non-limb fibroblasts (Fig. 1J). These results suggest that the pool of the candidate factors can convert non-limb bud fibroblasts to a state with at least some similarities to limb progenitors.

### Combinatorial overexpression of Prdm16, Zbtb16 and Lin28a induces limb progenitor marker expression in non-limb fibroblasts

Next, to identify which of the factors in our initial pool were responsible for the induction of limb progenitor marker genes, we examined the effect of withdrawing individual factors from the mix on the activation of the *Prx1* promoter, as reflected by GFP expression (18-1 factor assay; Fig. S5). Efficiency of the induction was measured as a GFP score, which was calculated by dividing the GFP positive area by total area staining with DAPI (Fig. 2A). We found that removal of any of 7 factors (Hoxd10, Zbtb16, Lhx2, Prdm16, Etv4, Tfap2a and Lin28a) resulted in a decrease in the GFP score, implying that these 7 factors were significant contributors to GFP induction (Fig. 2A). The combination of these 7 genes alone produced GFP positive cells efficiently, whereas withdrawal individual factors from the 7 factors pool decreased GFP scores (7-1 factor assay; Fig. S6A). We further conducted a 7-2 factor assay, in which combination of two factors were excluded from the 7 factors pool (Fig. S6B). We found that in both the 7-1 and 7-2 assays, Lin28a was necessary to yield a high GFP score (Fig. S6A and S6B). Moreover, Lin28 is required for induction of a second limb progenitor marker, Sall4 (Fig. S6A). Consistent with these results, Lin28 alone was sufficient to generate *Prx*GFP and Sall4 positive cell aggregates from non-limb fibroblasts (Fig. S6C), although other limb makers such as Lhx2 were not induced. To attain more complete reprogramming, we built on the Lin28a finding as a core factor, utilizing a Lin28a plus one factor assay (Fig. 2B, C). Although overexpression of Lin28a could not trigger Lhx2 expression (Fig. S6C), combination of Lin28a and either Zbtb16 or Prdm16 induced Lhx2 in addition to GFP and Sall4 (Fig. 2B, C). Combinatorial overexpression of both Prdm16 and Zbtb16 with Lin28a yielded even higher GFP scores (17.9 in Fig. 2B). Furthermore, transcript levels of representative limb progenitor genes were upregulated in the GFP-positive reprogrammed cells (Fig. 2D). Therefore, we defined these three as our core set of factors for limb reprogramming.

**Figure 2.**
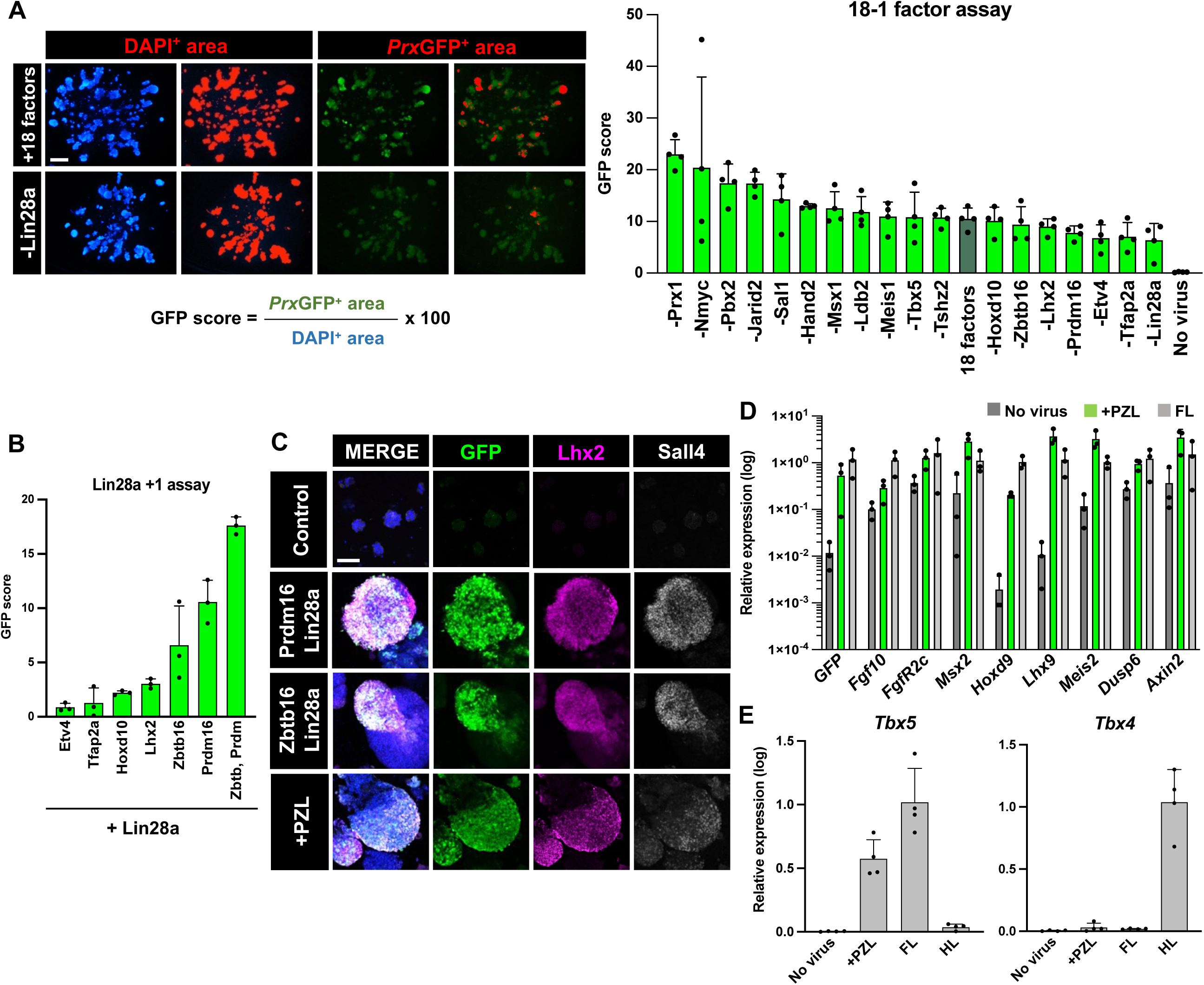
Identification of a minimal set of the reprogramming factors essential for imparting limb progenitor like-properties on non-limb fibroblasts. (A) Efficiency of *Prx*GFP induction was estimated as a GFP score by measuring GFP positive area per DAPI area. In 18-1 factor assay, each factor was withdrawn from the pools one by one (n = 4 gels each; see also Fig. S5). GFP score for the 18 factor-group was 10.57. Seven factors (Hoxd10, Zbtb16, Lhx2, Prdm16, Etv4, Tfap2a and Lin28a) that contributed to *Prx1*-GFP induction were tested for further screening as described in Fig. S6. The measured DAPI- or *Prx1*-GFP-positive area was pseudocolored in red. (B, C) GFP scores of Lin28a+1 factor assay. Combination of Lin28a with Prdm16, Zbtb16 or both (+PZL) yielded the highest GFP score and induced Lhx2 (magenta) and Sall4 (white) as well as *Prx*GFP (green) (n = 3 each). (D, E) qPCR for LP markers using controls (No virus), cells reprogrammed by overexpression of PZL, and LPs from E9.5 *Prx1*-GFP reporter embryos (n = 3 each in D, n = 4 each in E). GFP-positive reprogrammed cells and LPs were FAC-sorted beforehand. Error bars represent SD. Scale bars, 100 μm in (C), 1mm in (A).

The reprogramming factors we identified are expressed in both endogenous forelimb and hindlimb buds. To ask whether the reprogrammed cells acquired forelimb or hindlimb-like identity, we examined the expression levels of *Tbx5* and *Tbx4,* genes responsible for specification of the forelimb and hindlimb, respectively (Rodriguez-Esteban et al., 1999). We found that *Tbx5*, but not *Tbx4*, is induced in the reprogrammed cells, suggesting that the non-limb fibroblasts obtained forelimb-like traits through the overexpression of the reprogramming factors (Fig. 2E).

As noted above, we found that the clusters of reprogrammed cells were morphologically reminiscent of endogenous limb progenitors. To more rigorously assess this impression, we used forward scatter profiling to measure cell size, via flow cytometry. As expected from direct observation, the values of the reprogrammed cells were smaller than those of non-limb fibroblasts, and in the similar range to authentic limb progenitors (Fig. S7A). We also quantified and compared the size of nuclei (DAPI^+^) in unreprogrammed fibroblasts with that in the reprogrammed cells, and found the area of DAPI^+^ was decreased after reprogramming (Fig. S7B), again similar to the measured DAPI area of limb progenitors. Together, the reprogrammed cells share transcriptional and morphological similarities with legitimate early limb progenitors, and henceforth are termed as reprogrammed limb progenitor-like cells, or rLPCs.

### Overexpression of Egr1 suppresses limb progenitor proliferation and induces precocious differentiation of chicken limb progenitors

The results described above suggest that Pdrm16, Zbtb16 and Lin28a can in concert, convert non-limb fibroblasts into rLPCs. Lin28a in particular was the most indispensable in our 7-1 and 7-2 assays. Accordingly, we further investigated the role of Lin28a in rLPC reprogramming, in order to gain a more mechanistic understanding of the processes. Potential insight into this question came from consideration of its function as an iPSC reprogramming factor. In that context, Lin28a acts to block production of the Let-7 microRNA. This is significant because the *let-7* target, Lin41 suppresses translation of *Egr1,* which in turn antagonizes upregulation of pluripotency genes. Thus, in the presence of Lin28a, Lin41 activity promotes iPSC reprogramming (Ecsedi and Grosshans, 2013). Of note, *let-7a* is present in the chick limb buds and its expression level is increased as limb outgrowth proceeds (Lancman et al., 2005), corresponding to downregulation of *Lin28a* expression (Yokoyama et al., 2008). *Lin41* mRNA is also expressed in the early chicken and mouse limb mesenchyme (Lancman et al., 2005; fig. S8A). Conversely, Egr1 is not expressed in E9.5 or 10.5 mouse limb progenitors, nor is it seen in the forelimb-forming region of HH15 chicken embryos (Fig. 3A, S8A). However, Egr1 is detectable in differentiating limb progenitors and tenocytes of E13.5 mouse forelimb buds (Fig. 3B and S8B). These observations are consistent with Lin28a inhibiting *let-7a* in early limb buds, thereby preventing degradation of *Lin41*, and hence maintaining a limb progenitor state. Egr1 is also expressed in the non-limb fibroblasts used for reprogramming (Fig. 3B), suggesting that Egr1 may act to promote differentiation in the absence of reprogramming, as previously described in the human dermal fibroblasts used for iPSC reprogramming (Worringer et al., 2013).

**Figure 3.**
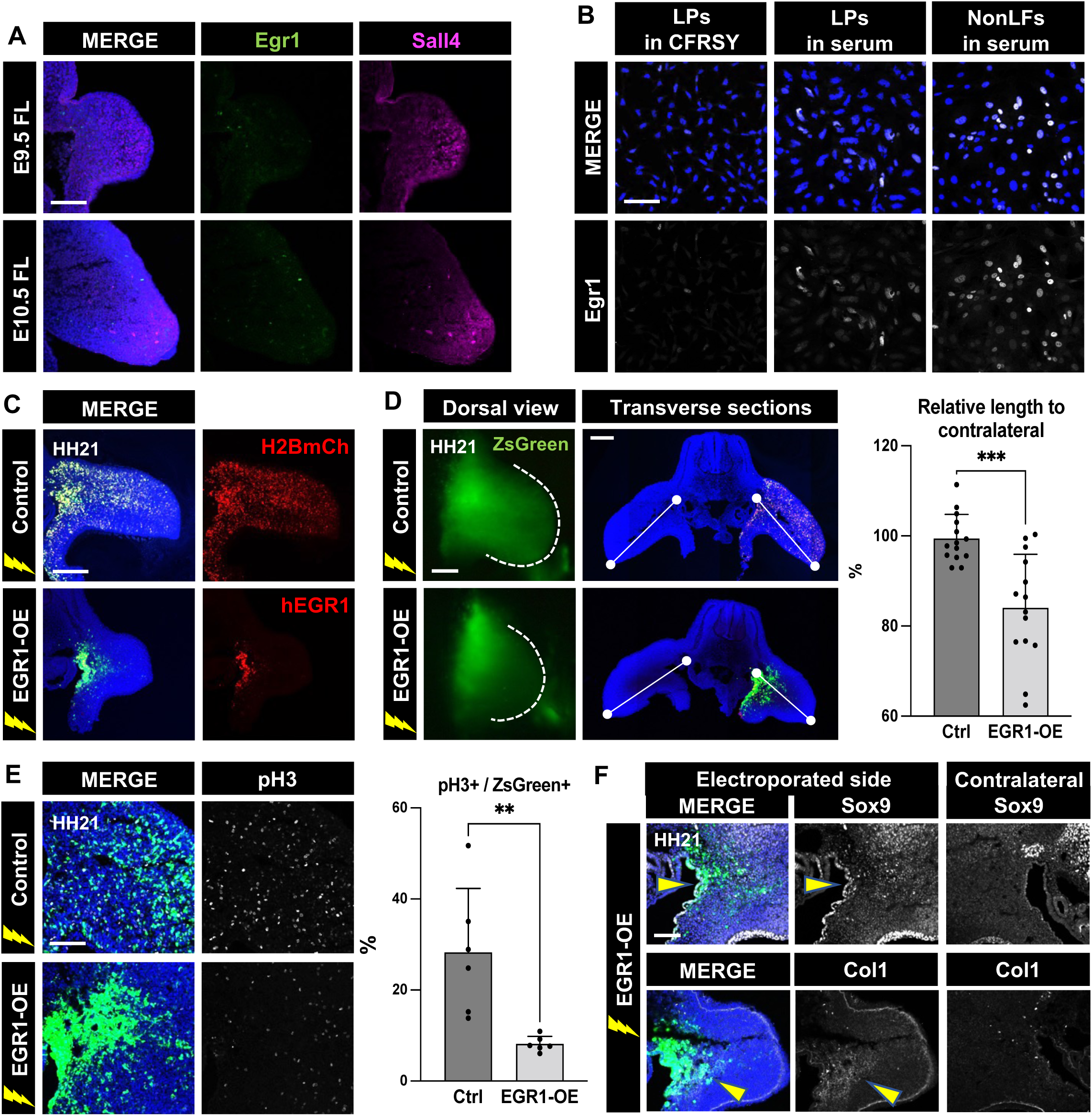
Misexpression of EGR1 disturbs limb bud outgrowth and induces precocious differentiation of limb progenitor. (A) Cross sections of E9.5 and E10.5 mouse FL buds stained with Egr1 (green) and Sall4 (magenta) antibodies. (B) E9.5 mouse LPs and NonLFs were cultured on petri dishes for 36 hrs in the presence of CFRSY or 10% FBS (serum), then were stained with an Egr1 antibody. (C) Plasmids carrying H2BmCherry-ires-ZsGreen1 (Control) or human EGR1-ires-ZsGreen1 (EGR1-OE) were electroporated into the chicken forelimb buds. Electroporated HH21 embryos were analyzed. (D) Overexpression of EGR1 inhibited lateral movement of limb mesenchyme. Relative length of the electroporated limbs to contralateral ones was measured (n = 14 limb buds each). (E) A mitotic marker phospho-Histone H3 (pH3) was detected by immunostaining in control and EGR1-electroporated limbs. pH3 positive cells per ZsGreen^+^ cells were counted (n = 6 each). (F) Immunostaining for Sox9 and Collagen I (Col1) in EGR1-electroporated or contralateral control limbs. \*\**p* < 0.01, \*\*\**p* < 0.001, a 2-tailed unpaired Student’s *t* test. Error bars represent SD. Scale bars, 100 μm in (A), (B), (E), 200 μm in (C), (D), (F).

To test if Egr1 indeed plays a role in the regulation of limb progenitors during limb development, human EGR1 coding sequences were electroporated into the somatopleural layer at the prospective forelimb level of HH13 chicken embryos, prior to the expression of endogenous *Egr1* mRNA (Fig. 3C-F). Limb mesenchyme electroporated with a control vector bicistronically expressing H2B-mCherry and ZsGreen was widely distributed in the limbs of HH21 embryos, whereas EGR1-transfected cells were located only around the coelomic epithelium, suggesting that overexpression of EGR1 either blocked these cells from entering the limb bud, or interfered with their distal migration (Fig. 3C, D). The EGR1-electroporated limbs were significantly reduced in length potentially attributable to the prohibition of limb progenitor migration, and also reflecting an attenuated level of cell proliferation, which was revealed by immunostaining for the mitotic marker phospho-Histone H3 (pH3), (Fig. 3D, E). Moreover, we found that the differentiation markers Sox9 and Col1 were induced in the EGR1-electroporated cells, meaning that these cells were precociously differentiated into chondrocytes or tenocytes (Fig. 3F). These data suggest that the EGR1 activity in limb progenitors drives cells towards differentiation, and hence its overexpression can disturb proper limb development, which may deteriorate the efficacy of rLPC reprogramming.

### Addition of Lin41 accelerates proliferation of rLPCs

Given that Egr1 appears to oppose the rLPC reprogramming (as previously observed for iPSC reprogramming) we decided to add Lin41 to the core set of reprogramming factors with the goal of further repressing expression of Egr1. Non-limb fibroblasts, carrying the GFP reporter under the control of the *Prx1* promotor were infected with lentivirus transducing Lin28a, Prdm16, and Zbtb16, with or without the addition of Lin41 (Fig. 4A). While we succeeded in converting non-limb fibroblasts into GFP^+^ putative rLPCs both in the presence and absence of Lin41(Fig. 4A and S9), the cell clusters that resulted from co-infection with Lin41 tended to be larger, and the proportion of pH3-positive cells was significantly higher, than in cultures reprogrammed without this factor (Fig. 4B). As expected, overexpression of Lin41 along with the other three reprogramming factors significantly decreased the number of Egr1 positive cells in comparison with non-limb fibroblasts and empty-virus infected controls (Fig. 4C). Moreover, cells reprogrammed with Lin41 expressed the same set of limb progenitor markers as cells reprogrammed by Lin28a, Prdm16, and Zbtb16 alone. (Fig. 4D-I and S9). These results suggest that the inclusion of Lin41 promotes cell proliferation of the rLPCs without adversely affecting the reprogramming process.

**Figure 4.**
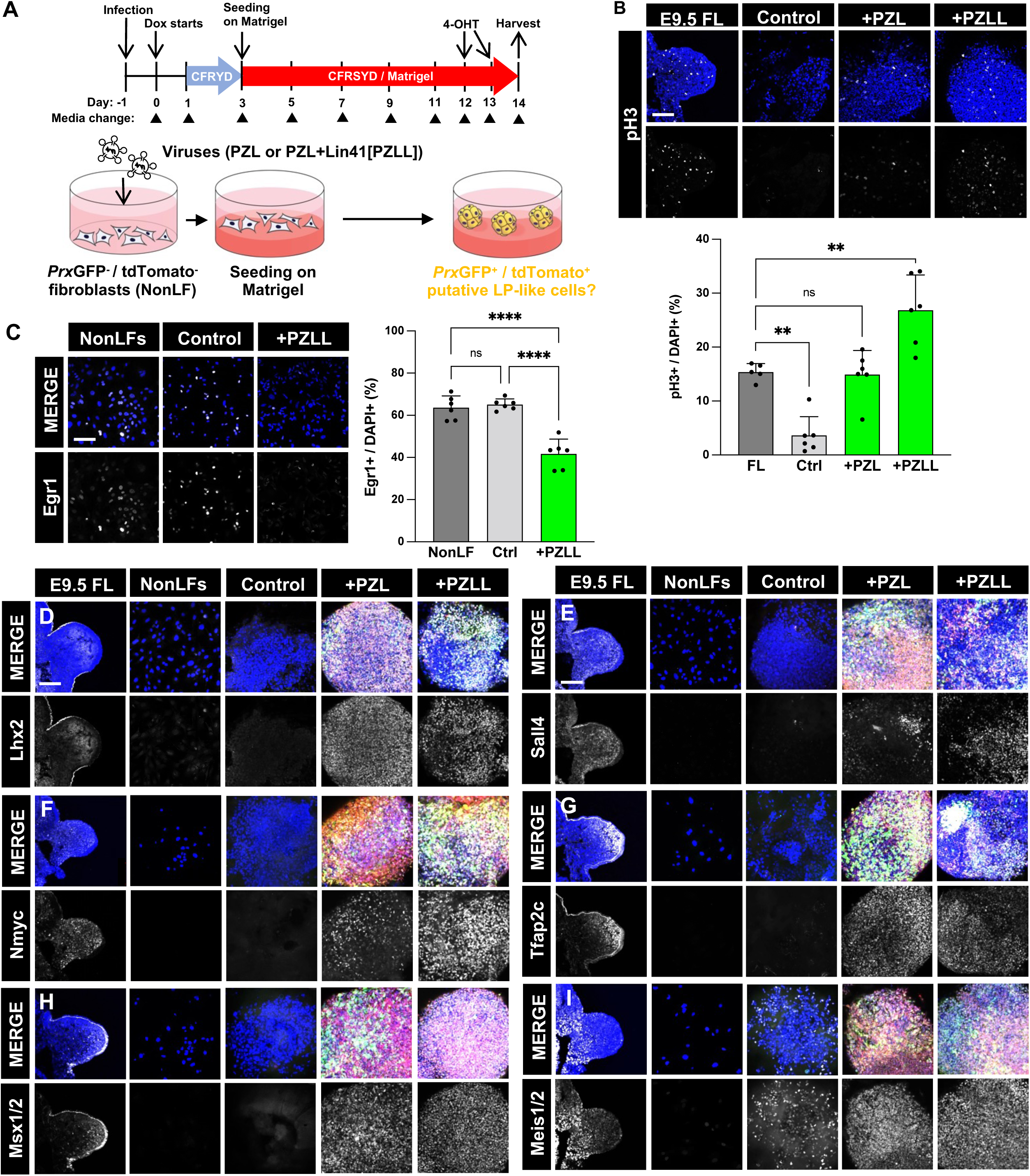
Addition of Lin41 to PZL stimulates proliferation of the rLPCs. (A) Schematics illustrating the modified reprogramming experiment. GFP/tdTomato-negative non-limb fibroblasts from *Prx1*-GFP/tdTomato reporter mice (Prx1-GFP-ires-CreER; CAG-LSL-tdTomato [Ai9]) were infected with tetO-lentiviruses carrying PZL and Lin41. Lentivirus carrying no transgene was used as Control. Doxycycline was administered during the culture. The cells overexpressing PZL or PZLL (PZL + Lin41) were seeded on Matrigel, and *Prx*GFP/tdTomato signals were examined at Day14. See also Fig. S9. (B) The number of pH3 signals was counted in E9.5 FL, Control, PZL- and PZLL-reprogrammed cells (n = 6 each). (C) Egr1 proteins were stained in NonLFs, Control and PZLL-reprogrammed cells. The number of Egr1 positive cells was quantified (n = 6 each). (D-I) LP markers were detected in the reprogrammed cells. E9.5 mouse FL and NonLFs were used as positive and negative control, respectively. In the MERGE panels for E9.5 FL and NonLFs, DAPI and signals for a target protein were merged. For Control, +PZL and +PZLL groups, DAPI, GFP, tdTomato and signals for the target were merged. Lhx2 (D), Sall4 (E), Nmyc (F), Tfap2c (G), Msx1/2 (H) and Meis1/2 (I) were induced in both PZL and PZLL reprogrammed cells. \*\**p* < 0.01, \*\*\*\**p* < 0.0001, one-way ANOVA. Error bars represent SD. Scale bars, 100 μm in (B-E).

### Reprogrammed rLPCs and primary limb progenitors share similar transcriptional profiles

Although the rLPCs that result from driving Prdm16, Zbtb16, Lin28a and Lin41 in non-limb fibroblasts show elevated expression of every early limb bud progenitor marker we tested, it was important to establish whether their global transcriptional profile approximated that of legitimate limb progenitors. To that end, we carried out a transcriptome-wide analysis by droplet-based single cell RNA sequencing (scRNAseq). Fibroblasts reprogrammed for 2, 4, 8 or 14 days (enriched for *Prx1*-GFP transgene expression by FACS, Fig. S10) were compared to E9.5 and E10.5 limb progenitors cultured *in vitro* under identical 3D matrigel conditions for 8 days. In addition, we assayed limb progenitors taken directly from E9.5, E10.0, E10.5 and E11.5/E12.5 stage embryos, as well as non-limb fibroblasts (cultured under either 2D or 3D conditions) as reference. In total, 74,268 single cell transcriptomes (Fig. S11, Table S1) were subject to dimensional reduction, low dimensional embedding (Brecht et al., 2018), graph-based clustering (Traag et al., 2019) and partition-based graph abstraction (PAGA) (Wolf et al., 2019).

The cells broadly cluster into seven distinct states, congruent with the different sources of the profiled cells (Fig. 5A, S12A). PAGA shows the relationship of these clusters to one another (Fig. 5A). At one end of this sequence is the cluster containing non-transfected non-limb fibroblasts cultured under 2-D conditions. Non-transfected non-limb fibroblasts (empty vector controls) placed into 3D culture are found in two adjacent clusters, shifted relative to the 2D cultured cells. In contrast, limb progenitors cultured under 3D conditions cluster separately from the non-limb fibroblasts. Limb progenitors taken directly from the embryo (ie. without being cultured *in vitro*) cluster separately from the 3D cultured progenitors, with distinct clusters for E9, E10, and E11 progenitors.

**Figure 5.**
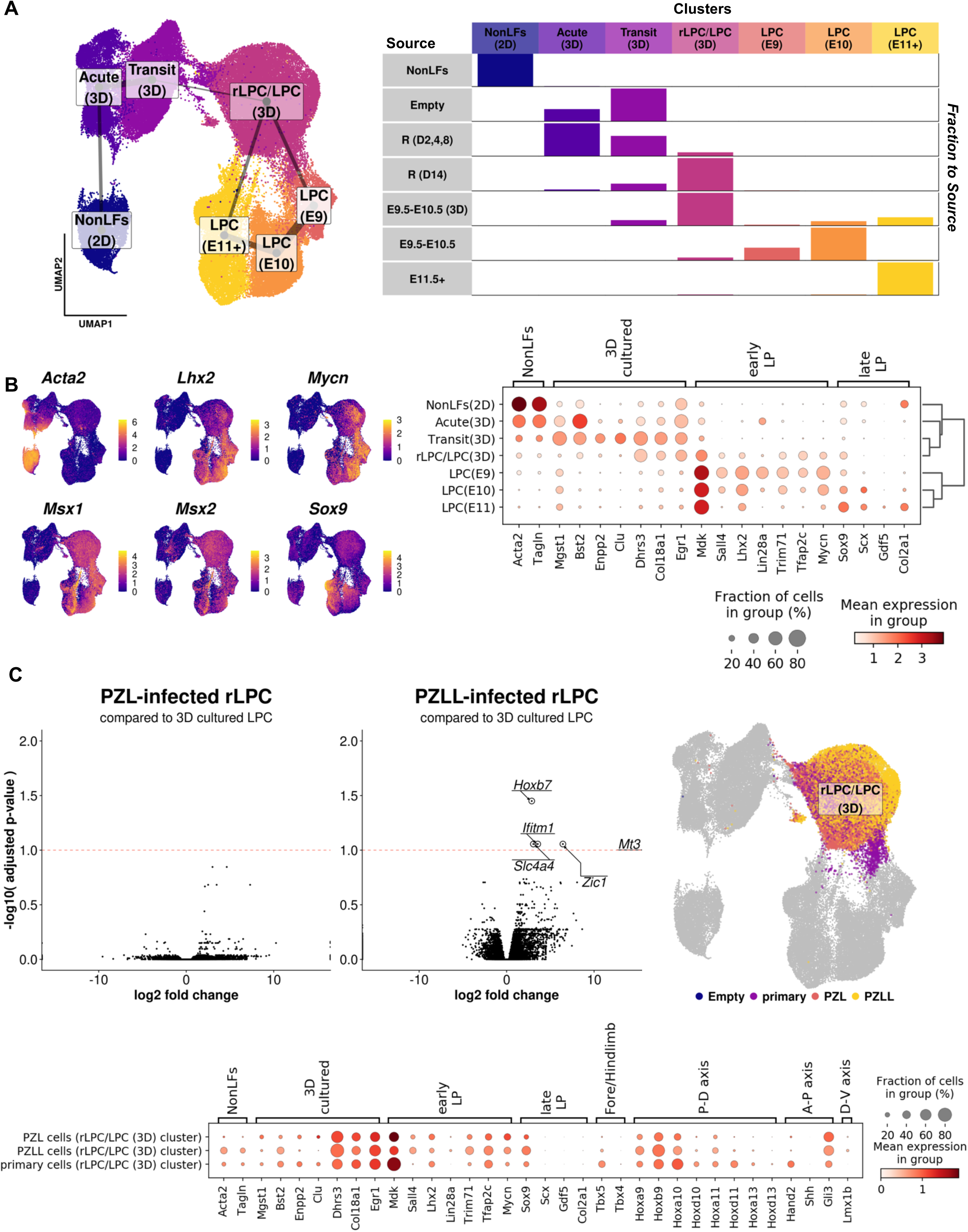
Single-cell RNA-seq analyses reveal global transcriptomic similarity between the rLPCs and endogenous limb progenitors. (A) Left panel: UMAP plot of NonLFs, limb progenitors (E9.5, E10.5, E11.5+), limb progenitors cultured for 8 days in matrigel culture (E9.5-E10.5 (3D)), cells infected with empty control virus (Empty) and reprogramming factors (R) sampled at different time points (D=days after culture). Overlaid are cluster labels by graph-based clustering (leiden, resolution=0.2), with edges between clusters from PAGA analysis. The thickness of edges represents the connectivity between clusters. Only the strong connection above threshold (0.05) were shown. Right panel: Split of cells by sample source and clusters. (B) Left panel: Expression of selected genes in UMAP coordinates. Right panel: Dot plot of select genes by clusters. (C) Volcano plot comparing PZL-infected/PZLL-infected rLPCs to 3D cultured LPCs, in rLPC/LPC (3D) cluster (Leiden, resolution=0.2). Adjustment of p-values were performed by pseudobulk aggregation of expression data by independent samples that were grown in 3D culture condition and comprise more than 100 cells for the rLPC/LPC (3D) cluster, using Benjamini-Hochberg adjustment (PZL: n=5, PZLL: n=3, primary: n=2). Red dotted line is threshold of adjusted p-value=0.1. Only comparison between 3D cultured LPCs and PZLL-infected cells have five genes above the threshold, circled and labeled. All genes more than 20 log fold changes, likely due to zero counts in one contrast, are put into infinity for better visualization. Right: UMAP plot showing the cluster and cells used for differentially expressed gene analysis in (C). Bottom panel: Dot plot of patterning genes in the rLPC/LPC (3D) cluster. All expression level in natural-log transformed UMI counts normalized by the total UMI counts per cell, maximal expression. PZL refers to Prdm16+Ztbt16+Lin28a (3-factor lentiviral expression). PZLL refers to Prdm16+Ztbt16+Lin28a+Lin41(Trim71) (4-factor lentiviral expression).

Most Non-limb fibroblasts subjected to reprogramming for 2, 4 or 8 days are found in the same clusters as control non-limb fibroblasts. Strikingly, however, the 3D cultured reprogrammed cells at day 14 completely overlapped with the cultured limb progenitors and were indistinguishable in terms of their transcriptome, showing essentially no differential gene expression and coverage (Fig. 5A, 5C and S13A). Moreover, this result was obtained whether the cells were reprogrammed with 3 or 4 factors (ie. Lin28a, Prdm16 and Zbtb16, with or without the addition of Lin41) (Fig. 5C); consistent with our finding (above) that Lin41 increases the proliferation of reprogrammed cells, but does not affect their differentiation state.

The UMAP pattern we observed can be further understood by reference to genes that characterize each cluster. Markers for non-limb fibroblasts (e.g. *Acta2*, *Tagln*) were quickly extinguished for all non-limb fibroblasts grown in 3D Matrigel culture, but only reprogrammed rLPCs upregulated markers similar to the early limb progenitors (eg. *Lhx2*, *Sall4*, *Tfap2c*, *Msx1/2*, *Mycn*). Notably, the reprogrammed cells did not upregulate markers of late-stage limb progenitors, such as *Sox9* (Fig. 5B). As noted above, the 3D cultured limb progenitors (and rLPCs) differ in their transcriptional profile from limb progenitors taken straight from the embryo. Genes differentially expressed by cells under these two conditions include targets of the signaling factors present in the culture media (Fig. S13B), and genes (such as ribosomal genes and cell cycle genes) reflecting the high proliferative state of reprogrammed cells *in vitro* (Fig. S11D, E, S13C).

While the 3D cultured limb progenitors fall into a single continuous cluster in this analysis, some distinctions can be observed within the clusters of limb progenitors directly taken from the embryo, reflecting differences in the patterning of the cells across the limb bud. Thus, there are subclusters representing Shh-expressing cells of the ZPA (zone of polarizing activity), and other genes indicative of cell variation across the anterior-posterior, and proximo-distal axes (Fig. S12D). In this context, the rLPCs, reprogrammed at day 14, mostly show expression of early proximal genes such as proximal *Hox* genes. In addition, the limb progenitors express either *Tbx5* or *Tbx4*, depending on their fore- or hindlimb origin, while rLPCs weakly express the forelimb marker *Tbx5*. Taken together, the transcriptome analysis suggests that the reprogrammed cells attain an early forelimb progenitor state, in an active state of proliferation, without evidence of late patterning or differentiation (Fig. 5D).

### Trajectory analysis reveals the sequence of events during the reprogramming of non-limb fibroblasts into limb progenitor-like cells

Having established that driving the expression of Lin28a, Prdm16, Zbtb16 and Lin41 indeed drives non-limb fibroblasts to a limb progenitor-like state, we wanted to better understand the process by which this occurs. Accordingly, to explore the transcriptional dynamics of the reprogramming, we sub-clustered the cells at higher resolution (Figure 6A, S14B) and turned to optimal-transport analysis (Waddington Optimal Transport, WOT) (Schiebinger et al. 2019). WOT infers the growth rates, and the ancestor-descendant relationship of cells across time points utilizing the transcriptome information of individual cells at intermediate time point samples (Fig. S14A). This in turn is used to construct probabilistic trajectories to specific fates (Fig. 6B, Fig. S14D).

**Figure 6.**
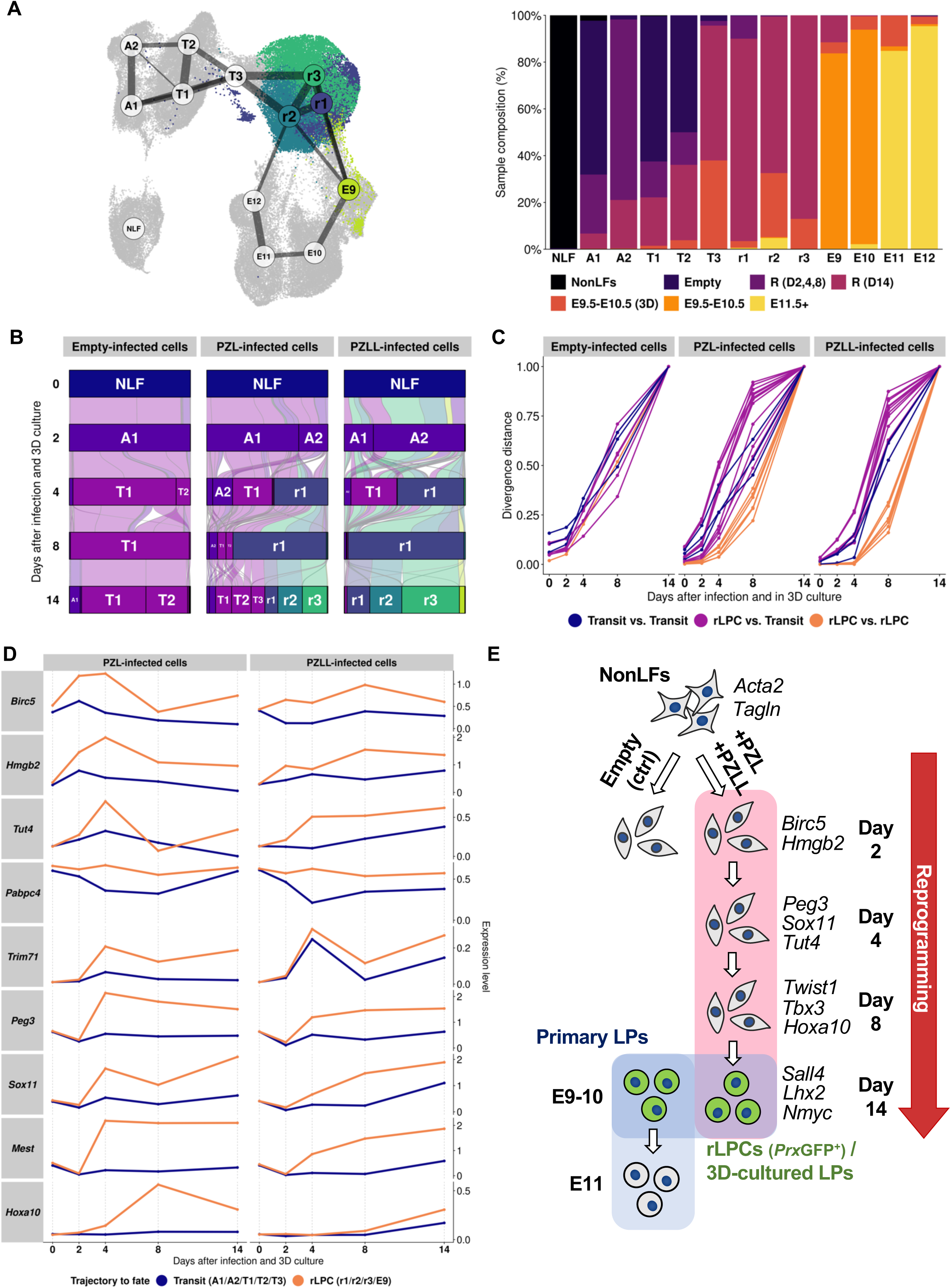
Optimal transport analysis delineates transitions of reprogramming of the rLPCs from non-limb fibroblasts. (A) Left panel: UMAP plot with fine clusters (Leiden clustering, resolution=0.4), overlaid with edges between clusters from PAGA analysis. The thickness of edges represents the connectivity between clusters. Only the strong connection above threshold (0.1) were shown for clarity. Right panel: the composition of each clusters according to the sample source in stacked column graph. The clusters are roughly ordered from the initial starting material (NonLFs) to the later stages of limb progenitor cells. (B) Alluvial (flow) plot based on the transition matrix inferred by Waddington Optimal Transport (WOT) analysis. WOT analysis generates temporal couplings between sets of cells between time points. The initial width of each alluvial segment represents the probability of transition of the group of cells from the earlier state to later state. The final width incorporates the estimated growth rate of the destination cell cluster. Thus, wider width than the initial starting point represent expansion (proliferation) after transition, whereas narrower width means contraction (cell death or stasis) to the next time point. All alluviums are colored by the final (Day 14) fate of the cells (See also Fig. S14D for individual highlights). (C) The fraction of transcriptional divergence accrued at intermediate time points between trajectories towards final fate. Each lines represent a comparison between two distinct trajectories, grouped and colored by the final fate of the two populations. (D) Changes of mean expression levels of individual genes at a given time point weighted by the probability of the final fate inferred by WOT. All expression level in natural-log transformed UMI counts normalized by the total UMI counts per cell. (E) Schematic diagram of reprogramming. PZL refers to Prdm16+Ztbt16+Lin28a (3-factor lentiviral expression). PZLL refers to Prdm16+Ztbt16+Lin28a+Lin41(Trim71) (4-factor lentiviral expression).

At 14 days after infection and 3D culture, the infected, 3D cultured cells are clustered into four rLPC sub-states (r1, r2, r3, and E9) as well as three transit sub-states (T1, T2, T3), which are used as fates to construct trajectories (Fig. 6A, B, Fig. S14, S15). The four rLPC substates are distinguished by the relative similarity to the E9.5 stage limb progenitors in vivo, where E9 cluster cells grouped together with early E9.5 limb progenitors, with r1/r3 clusters neighboring to the E9 cluster, and r2 cluster close to both E9 and a subset of *Osr1*+ E12.5 limb progenitors (Fig. S14C). Moreover, the r1 population arises as early as Day 4 after infection and 3D culture, with strong proliferative signature (Fig. S14B, C) whereas r2, r3, and E9 populations are only detected by Day 14. On the other hand, both the acute-phase (A1, A2) as well as transit (T1, T2, T3) clusters display markers of various inflammatory markers, with the A1, A2 cluster showing high expression of the transgene *Lin28a* (Fig. S14C).

The reconstructed rLPC trajectories suggest that by Day 4, the r1 cluster with high proliferative activity arise that dominate the contribution to the subsequent successful reprogrammed state (Fig. 6B, S14D). Comparing the transcriptional divergence between the trajectories, the trajectories leading to rLPC states remain close each other until Day 8, whereas they all quickly diverge from others, suggesting that successful reprogramming is determined at early phases of infection and culture and those in the successful trajectory remain plastic to a particular rLPC fate (Fig. 6C). Moreover, the reconstructed trajectories provide differentially expressed genes at early time points that are associated to the successful rLPC fate (Fig. 6D, S15A). At acute infection phase, genes countering apoptosis and promoting proliferation are found to be upregulated. Interestingly, the initial level of lentiviral expression as assessed by the counts of Woodchuck Hepatitis Virus Posttranscriptional Response element (WPRE) reads appear to be negatively associated with the rLPC trajectory from others, suggesting that expression of the transgenes was downregulated during the latter process of the reprogramming and may not be required for rLPC production. It is followed by the endogenous upregulation of *Lin41* (*Trim71*) as well as genes involved in mRNA stability (*Tut4*, *Pabpc4*) and transcription factors *Peg3* and *Sox11* that distinguish the successful rLPC trajectories from others. This is true for cells reprogrammed with either 3 or 4 factors (Fig. 6D, Fig. S15A). Lastly, transcription factors involved in patterning appear later at Day 8. Other factors, such as *Prdm16* as well as *Zbtb16* were found to be differentially expressed at later phases in reprogramming.

### Reprogrammed rLPCs differentiate into limb cell types and respond properly to limb patterning cues in vitro

While rLPCs closely resembled limb progenitors at a transcriptional level, it was important to also establish whether they were capable of behaving as such at a functional level. To that end, we first asked if they acquired the capability to differentiate into cell types normally found in the developing limb bud. In this instance reprogramming was done without Lin41, as we wanted the rLPCs to be able to freely differentiate once culture conditions were changed. After reprogramming, GFP positive rLPCs were sorted by FACS and cultured in 96 well plastic plates under micromass culture conditions (a well-established *in vitro* system, used to study the differentiation of limb progenitors) in the presence of the growth factors we optimized for keeping limb progenitors undifferentiated. When the cultures became confluent, the growth factors were withdrawn to promote differentiation of the cells, and they were grown for 8 additional days. The chondrogenic capacity of the cells was then analyzed by Sox9 protein and Alcian blue staining, and qPCR for Sox9 (an early chondroprogenitor marker) and *Aggrecan1 (Agc1)* (a mature chondrocyte marker). We also assessed the capacity to differentiate into connective tissue by looking at expression levels of *Scleraxis* (*Scx*), a marker for tendon and ligament precursors (Schweitzer et al., 2001), and *Odd-skipped related 2* (*Osr2*) a gene known to be required for specification of joint cells (Gao et al., 2011). Multiple clusters of differentiated reprogrammed cells stained positively with Sox9 and Alcian blue whereas unreprogrammed non-limb fibroblasts did not (Fig. 7A, B). Additionally, transcript levels of *Sox9* and *Agc1* were upregulated in the differentiated reprogrammed cell cultures, indicating that the rLPCs have acquired chondrogenic potential (Fig. 7C). Moreover, the level of expression of *Scx* and *Osr2* in these differentiated reprogrammed cells was increased (Fig. 7C), indicating that the reprogrammed cells are capable of differentiating into connective tissue cell types as well.

**Figure 7.**
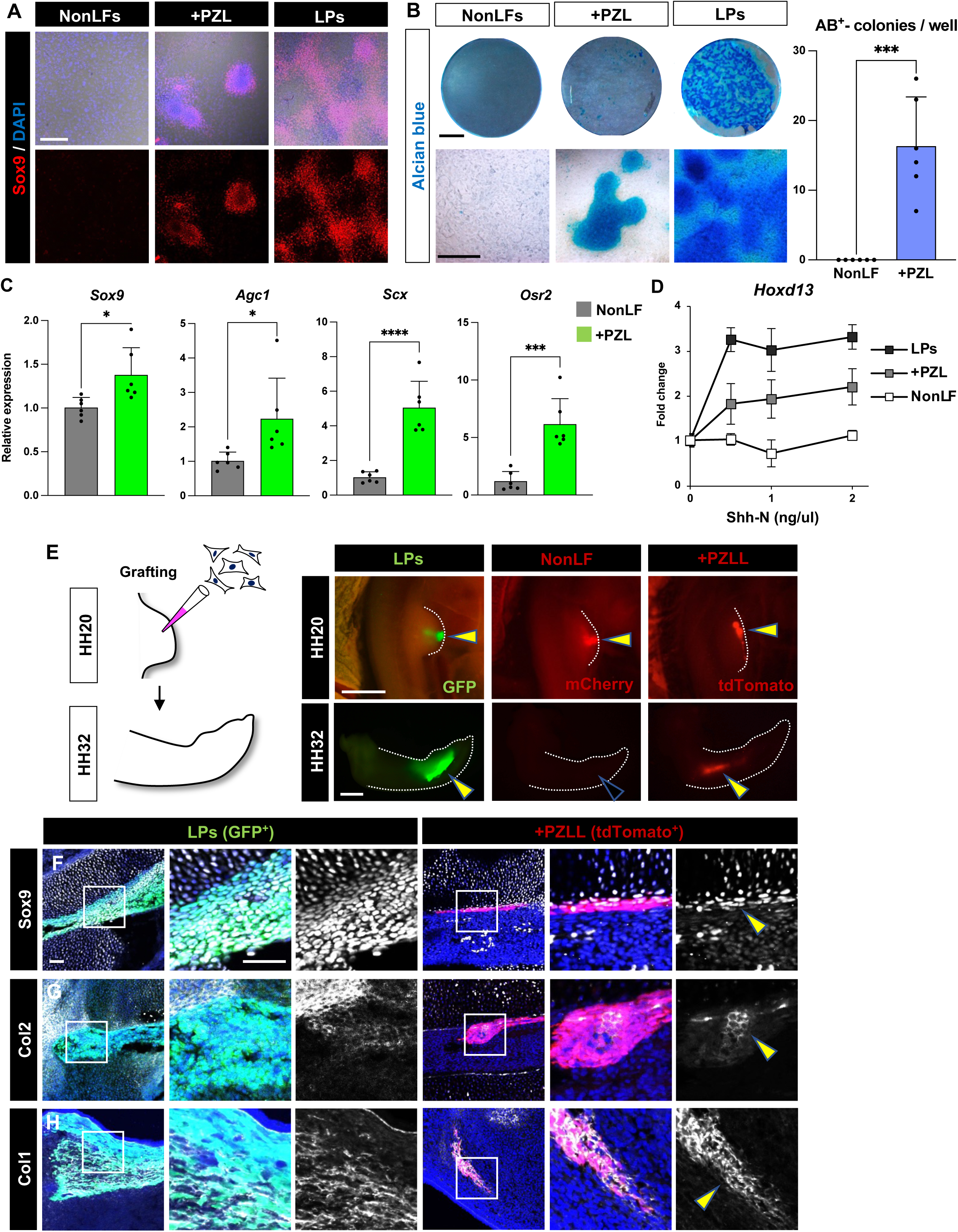
The rLPCs exhibit differentiation potency towards chondrocytes and tenocytes. (A, B) Micromass cultures to test *in vitro* chondrogenesis capacity of the reprogrammed cells. Sox9 or Alcian blue positive clusters emerged from the reprogrammed cells. The number of Alcian blue positive clusters in NonLFs and the reprogrammed cell groups were counted (n = 6 wells for each). (C) qPCR analyses for *Sox9*, *Aggrecan1* (*Agc1*), *Scleraxis* (*Scx*) and *Osr2* (n = 6 each). FL cells from E9.5 *Prx1*-GFP embryos were micromass-cultured as well and used as positive controls. (D) Shh ligand and *Hoxd13* gene expression titration curves. Samples were treated for 24 hrs with varying levels of Shh ligand (0, 0.5, 1 and 2 ng/μl; n = 3 for each group and time point). (E) E9.5 CAG-GFP mouse LPs, NonLFs expressing mCherry and FAC-sorted tdTomato PZLL-reprogrammed cells were transplanted into HH20 (E3.5) chick FL buds. 4 days after the grafting, the limbs were harvested at HH32 (E7.5). The grafted GFP-LPs and tdTomato-reprogrammed cells were seen in the HH32 limbs (yellow arrowheads), while mCherry-NonLFs were not detectable (a black arrowhead). (F-H) The harvested HH32 limbs were sectioned and stained with Sox9 (F), Collagen II (Col2, G) and Col1 (H) antibodies. A fraction of the grafted LPs (n = 7) and tdTomato-reprogrammed cells marked by yellow arrowheads (n = 3) were positive for each marker. \**p* < 0.05, \*\**p* < 0.01, \*\*\**p* < 0.001, \*\*\*\**p* < 0.0001, a 2-tailed unpaired Student’s *t* test. Error bars represent SD. Scale bars, 100 μm in (F), 1 mm in (A), (B, the lower bar), (E), 2mm in (B, the upper bar).

We next asked whether the reprogrammed cells would respond to patterning signals in a manner similar to endogenous limb progenitors. The optimized media we established for maintaining limb progenitors in culture already contained RA and Fgf8, two signals important for the establishment of proximodistal patterning in the limb buds (Cooper et al., 2011). We therefore examined targets of each of these factors that are up-regulated during the normal patterning of the developing limb bud. *Meis2*, a downstream effector of RA signaling in the proximal limb bud and *Dusp6*, a readout of Fgf signaling in the distal limb bud, were both activated in the reprogrammed cells (Fig. 2D). A third important morphogen in the early limb bud is Sonic hedgehog (Shh), a polarizing signal acting along the anterior-posterior limb axis. To examine response to Shh, we assayed the induction of Hoxd13, a key target in the limb bud (Tarchini et al., 2006, Rodrigues et al., 2017). After 24 hours of exposure to Shh, *Hoxd13* upregulation was observed in a dose-dependent manner in both reprogrammed cells and legitimate limb progenitors whereas it was not seen in unreprogrammed non-limb fibroblasts (Fig. 7D). Taken together, the rLPCs appear to have differentiation and patterning potential in vitro similar to those exhibited by endogenous limb progenitors.

### Reprogrammed rLPCs differentiate into limb cell types in vivo

While these results indicate that rLPCs can respond similarly to limb progenitors under artificial conditions *in vitro*, and generate limb–specific cell types in that setting, it was important to determine whether they could also integrate into a developing limb bud and differentiate appropriately *in vivo*. To test this, we exploited a tetracycline-inducible lentivirus system (Stadtfeld et. al., 2008) (Fig. 4A), so that the reprogramming factors would be under temporal control *in vitro*, and would be inactivated upon transplantation *in vivo*. We also needed to be able to follow the transplanted cells as they differentiated, even if they ceased to express GFP from the *Prx1* promoter. To that end, we harvested non-limb fibroblasts from mouse embryos carrying a dual reporter. One transgene (*Prx1-*CreER-IRES-GFP) expresses both CreER and GFP in limb progenitors. The GFP activity is therefore lost when the cells differentiate into a state that no longer drives expression from the *Prx1* promoter. However, a second transgene (R26-CAG-LSL-tdTomato) is irreversibly activated in any cell even transiently expressing CreER in the presence of tamoxifen (Fig. 4A and S9). Thus, derivatives of rLPCs will be marked as red, regardless of whether or not they continue to express GFP from the *Prx1* promoter.

rLPCs were generated by introducing Lin28a, Prdm16, Zbtb16 and Lin41 to non-limb fibroblasts via the lentivirus vectors, cultured in the presence of doxycycline, as well as the factors optimized for maintaining limb progenitors (Fig. 4A). These reprogrammed cells were then cultured for 2 days without doxycycline, or the other limb progenitor-maintenance factors, and then the cells were xenografted into limb buds of HH20 chicken embryos (Fig. 7E). Despite heterospecific transplantation, grafted authentic limb progenitors derived from E9.5 CAGGS-GFP mice readily integrated into chicken wing buds, as previously reported (Fig. 7E) (Izpisua Belmonte et al. 1992). By contrast, almost all mCherry-transfected non-limb fibroblasts were eliminated from the chicken limbs 4 days after they were grafted (Fig. 7E). Similar to the endogenous limb progenitors, the grafted reprogrammed cells stayed within the host limbs over this time period (Fig. 7E). Strikingly, subsets of the tdTomato-positive reprogrammed cells were seen to differentiate into chondrocytes marked by Sox9 or Col2al, and into tenocytes that were stained with an antibody against Col1, similar to legitimate mouse Limb progenitors transplanted into the chicken limbs (Fig. 7F-G). Thus, we conclude that reprogrammed cells are multipotent, are able to participate in limb development, and can generate normal limb tissues *in vivo*.

### Reprogramming human fibroblasts into cells resembling limb progenitors

The identification of a set of genes capable of reprogramming embryonic mouse non-limb fibroblasts into rLPCs holds the promise of providing new insight into the specification of the limb bud. In addition, however, this work suggests a potential route towards providing cells that can be used in a therapeutic setting, provided the process can be replicated starting with adult human cells. While a full characterization of human rLPCs would be beyond the scope of this study, we wanted to at least get an indication of whether the reprogramming factors we identified in the murine system would have a similar effect in human fibroblasts. To that end, adult human dermal fibroblasts were infected with lentiviruses transducing our three core reprogramming factors, Lin28a, Pdrm16 and Zbtb16, and were then placed in 3D culture under limb progenitor maintenance conditions. After 18 days, cell aggregates emerged, resembling plated mouse limb bud cells as well as those seen when reprogramming mouse non-limb fibroblasts (Fig. S16A). We examined the expression of several limb progenitor markers (SALL4, LHX2 and NMYC) as well as EGR1 in these cells. All three limb progenitor markers were up-regulated in comparison with control human dermal fibroblasts, while EGR1 expression was diminished (Fig. S16B, C). Of note, the expression patterns of NMYC and EGR1 were mutually exclusive (Fig. S16C).

To get a more complete understanding of the transcriptional changes resulting from the reprogramming of the human dermal fibroblasts, we undertook a single-cell transcriptomic analysis of the human cultures infected with the Lin28a, Pdrm16, Zbtb16 lentiviruses, with or without co-infection of Lin41. Cells cultured in the 3D limb progenitor maintenance conditions for 18 days were compared to control human dermal fibroblasts grown in the same conditions (Fig. S17A). These data further support the down-regulation of dermal fibroblast markers and up-regulations of limb progenitor markers (Fig. S17B). A limitation of using human cells is the lack of legitimate embryonic human limb progenitors for comparison. Therefore, the human reprogrammed and control samples were aligned with the mouse single cell transcriptome embedding. This analysis indicates that the reprogrammed human dermal fibroblasts aligned with the early mouse limb progenitor state (Fig. S17C).

Finally, to get preliminary indication of whether the reprogrammed human rLPCs have some of the same differentiation potential as limb bud cells, we conducted xenograft experiments in which the dissociated putative reprogrammed cells were transplanted into chicken limb buds. Unlike mouse non-limb fibroblasts, the grafted human dermal fibroblasts were able to engraft in the chicken limbs, however, they were completely excluded from cartilage elements and showed no Sox9 expression (Fig. S16D). By contrast, a fraction of the grafted reprogrammed cells integrated into Sox9^+^ cartilage (Fig. S16D), implying that the cells could differentiate into chondrocytes. The percentage of transplanted cells incorporated into the cartilage seemed to be much lower than with the mouse rLPCs. However, that was to be expected as, unlike the transgenic mouse cells, human dermal fibroblasts lacked the *Prx1*-GFP reporter, and hence the cultures could not be enriched for reprogrammed cells by FACS prior to transplantation. Taken together, these results suggest that human dermal fibroblasts are indeed transformed by the same reprogramming factors as in the mouse, towards a state that at least has characteristics in common with limb progenitors.

## Discussion

In this study, we have established long-term culture conditions to maintain limb progenitors, identified factors that are sufficient to reprogram non-limb fibroblasts into rLPCs, and validated their similarity to limb progenitors via multiple criteria.

### Optimized 3D culture conditions for long-term maintenance of limb progenitors

Identifying adequate culture conditions for maintaining stem cells being targeted is known to have been a key factor in the success of other reprogramming studies. For instance, the Yamanaka factors failed to reprogram mouse embryonic fibroblasts to iPSCs in the absence of leukemia inhibitory factor (LIF) and feeder cells (Takahashi and Yamanaka, 2006). Since our previous culture condition for limb progenitors (Cooper et al. 2011) was effective only for the short term, we sought to optimize the conditions for long-term maintenance of limb progenitors. Ultimately, we found that a cocktail of CHIR90021 (a GSK3β antagonist) Fgf8, RA, SB431542 (a Bmp/TGFß inhibitor) and Y-27632 (a Rock inhibitor) will maintain limb progenitors in a HA or Matrigel 3-D matrix for an extended period of culture. Although RA is necessary to keep cells in the progenitor state through activation of limb progenitor genes such as *Meis1/2* and by blocking chondrogenic differentiation (Cooper et al., 2011), RA can also induce apoptosis as seen in interdigital mesenchyme. The RA-induced apoptosis is partially mediated by Bmp7 (Dupé et al., 1999), thus TGFβ/BMP antagonist SB431542 may not only inhibit differentiation of limb progenitors but also block cell death during culture. In addition, it is noteworthy that the endogenous RA concentration is higher in the anterior part of the embryo than that in the posterior region and thereby promotes induction of *Tbx5*, but not *Tbx4*, during forelimb initiation (Nishimoto et al., 2015). It is therefore likely that RA also contributes to upregulation of Tbx5 in rLPCs during reprogramming, and is thus responsible for the forelimb-like characteristics of these cells.

### Possible roles of the reprogramming factors

Given that overexpression of Lin28a alone is capable of inducing *Prx*GFP and Sall4, we consider Lin28a as a central reprogramming factor. By contrast, exogenous Tbx5 and Nmyc were dispensable for rLPC reprogramming despite their necessity for normal mouse limb development (Agarwal et al., 2003). Intriguingly, *Lin28, Sall4, Nmyc, Tbx5* and *Lin41*, mRNAs that are transcribed in early limb progenitors, are suppressed by members of the *let-7* miRNA family in other contexts, including the regulation of embryonic stem cells, iPSC reprogramming, and during cardiogenesis (Wang et al., 2013). Thus, there is a possibility that Lin28a indirectly upregulates expression of limb progenitor-specific genes globally, by blocking *let7* miRNA activity, thereby triggering rLPC reprogramming. We also find that Lin41 promotes mouse rLPC proliferation and maintenance in a progenitor state. In our scRNAseq analysis, endogenous Lin41 upregulation was an early gene expression signature at the time when highly proliferative r1 subpopulation arise, and a lower level of endogenous Lin41 expression at later time points in a subset of rLPCs lacking Lin41 overexpression were associated with rLPC subpopulation which showed transcriptomic similarity to later phase limb bud cells (r2), whereas rLPC trajectories that maintained high level of Lin41 expression resulted in rLPC fates that showed transcriptional similarity to early limb bud cells. A similar result was seen with reprogrammed human dermal fibroblasts. In scRNAseq analysis, Lin41-overexpressing cells were partially aligned with E9.5 mouse limb progenitors, whereas reprogrammed cells without Lin41 were separated from the early limb progenitors, suggesting a role for Lin41 in keeping reprogrammed cells in the undifferentiated early limb progenitor state. Mechanistically, Lin41 is likely to inhibit translation of Egr1, but not mRNA transcription, given that transcript levels of *Egr1* are not decreased in Day 14 mouse and Day 18 human reprogrammed cells according to our scRNAseq analysis. Lin41 is also known to ubiquitinate the tumor suppressor p53 in murine embryonic stem cells, thereby antagonizing cell death and differentiation pathways (Nguyen et al., 2017). As suppression of p53 promotes iPSC reprogramming (Kawamura et al., 2009), perhaps Lin41 potentiates rLPC reprogramming through its ubiquitinase activity. This raises the possibility that there is a“*let7* barrier” that may hamper rLPC reprogramming as seen in iPSC reprogramming (Worringer et al., 2013). In that context, *let-7* miRNAs suppress stemness factors including *Oct4*, *Nanog*, *Sox2* (Melton et al., 2010), and *Myc* and *Lin41* (Worringer et al., 2013).

In concert with Lin28a, Prdm16 and Zbtb16 are each capable of inducing Lhx2 expression in non-limb fibroblasts. The role of Prdm16 in limb development has not been previously characterized. Prdm16 contains protein interacting zinc-finger and histone lysine methyltransferase domains and is known as a crucial regulator of adipose development, with implications for several processes including energy homeostasis and glucose metabolism (Chi and Cohen, 2016). Considering that accelerated metabolism is a key driver for iPSC reprogramming and tumorigenesis, and rapid proliferation is one of the hallmarks of early limb progenitors (Spyrou et al., 2019), Prdm16 may contribute to rLPC reprogramming by enhancing the metabolic status of non-limb fibroblasts in addition to inducing limb progenitor-specific genes such as Lhx2. Unlike Prdm16, the involvement of Zbtb16 in limb development has been described previously. Zbtb16, which is also a zinc-finger transcription factor, regulates the expression of several *Hox* genes, including *Hox10,* downstream of Sall4, and is required for proximal development of the mouse limb (Barna et al., 2000). Whether Zbtb16 similarly controls *Hox* expression during rLPC reprogramming is a topic for future investigation.

### Potential of rLPCs for clinical application

As rLPCs have the potential to differentate into chondrocytes and connective tissues, rLPCs could, in principle, be harnessed for regenerative therapies in the future. Previously, endogenous limb progenitors and iPSC-derived limb progenitor-like cells have been shown to enhance regenerative processes when transplanted into amputated frog limbs and mouse digit tips, respectively (Lin et al., 2013; Chen et al., 2017). 3D spheroids of limb progenitor-like cells also can be induced from mouse embryonic stem cells (Mori et al., 2019). None of these studies, however, including our own, have demonstrated that induced or reprogrammed limb progenitors have the capacity, on their own, to give rise to a limb-like structure, patterned along various axes and containing appropriate differentiated tisssues. In principle, this can be tested by constructing a “recombinant limb”, in which dissociated limb mesenchyme (or, in principle, rLPCs) are pelleted, and packed into an empty shell of limb ectoderm, and grafted onto a host embryo (Zwilling, 1964, Ros et, al., 1994). Such recombinant limbs made with limb progenitors make well formed limb-like structures. However, as the recombaint limb assay is only feasible with avian embryos, a recombinant system using reprogrammed avian cells will be required.

Our study may also open the way to *in vivo* direct rLPC reprogramming (Zhou et al., 2008). By overexpressing the reprogramming factors in dermal fibroblasts at an amputation site of a human limb, cells might be reprogrammed towards a limb progenitor state, thereby potentiating the *in situ* development of a limb-like structure. Of note, two of the reprogramming factors, *Lin28* and *Prdm16* are re-expressed in blastema of regenerating appendages in other systems (Rao et. al., 2009; Yoshida et. al., 2020). While such therapeutic applications will require a great deal of further work, the study described here provides a more immediate platform for interrogating the molecular control of the limb progenitor state.

## Acknowledgements

We thank Drs. Gufa Lin (University of Minnesota), Yasu Kawakami (University of Minnesota), Johanna Kowalko (Florida Atlantic University), Jessica L. Whited (Harvard Medical School) and Daisuke Saito (Kyushu University) for helpful discussions. We also thank the Single Cell Core (Harvard Medical School), the Flow Cytometry Core (Brigham and Women’s Hospital), and the Biopolymers Facility (Harvard Medical School) for providing experimental platforms used in this work. This work was supported by NIH grant HD03443 (to C.J.T.). Y.A. was a recipient of fellowships from the Naito foundation and JSPS for research abroad. E.L. was a recipient of a fellowship from the NSF 1612264.

## Author Contributions

Y.A., C.L., A.R. R., C. C., R. R. T. and E. G. L. conducted the experiments; Y. A., C. L., A. R. R., C. C., R. R. T., P. T., D. C and J. G. analyzed the data; J. P. V. contributed new reagents; C.E.S., J.G.S., O.P., and C.J.T. supervised the work; Y.A., C.L., and C.J.T. wrote the first draft; and all authors revised the manuscript.

## Declaration of Interests

The authors declare no competing financial interests.

**Supplemental figure 1**

**Optimization of conditions for culturing endogenous mouse limb progenitors,**

**Related to Figure 1**

(A) Limb progenitors (LPs) from forelimbs (FL) of E9.5 CAG-GFP mouse embryos were cultured in either 10% FBS/DMEM (Serum condition) or media supplemented with Chir99021 (3 μM), Fgf8 (150 ng/ml), Retinoic acid (25 nM) (CFR condition) for 4 days. (B) LPs from E9.5 *Prx1*-CreER-ires-GFP mouse embryos were dissected out and cultured for 10 days under Serum, CFR and CFRSY (CFR plus Y-27632 and SB431542) conditions. The cells were stained by using GFP (green), Lhx2 (magenta) and Sall4 (white) antibodies. Scale bars, 100 μm in (B), 200 μm in (A).

**Supplemental figure 2**

**Chicken limb progenitors cultured in CFRSY/HA-gel condition maintain differentiation potentials into chondrocytes and tenocytes, Related to** **Figure 1**

(A) LPs from HH18 GFP chicken embryos were cultured in hyaluronan (HA)-based hydrogels, in the presence of CFRSY for 8 days, and then they were dissociated and transplanted into HH20 chick FL buds. The grafted limbs were harvested at HH32, 4 days after transplantation, and sectioned followed by staining for Sox9. The grafted cells were seen in Sox9-positive cartilage (yellow arrowheads). (B) The grafted GFP cells were stained with Collagen I antibody (yellow arrowheads). (C) The cells were MHC, a muscle marker, negative (black arrowSheads), but closely associated with MHC-positive muscles (red). Scale bars, 100 μm in (A-C).

**Supplemental figure 3**

**Expression of limb progenitor marker genes is maintained in mouse LPs cultured under CFRSY/Matrigel condition, Related to** **Figure 1**

(A) *Prx*GFP^+^ LPs were cultured on 50% Matrigel in media supplemented with CFRSY for 8 days. *Prx*GFP, Lhx2 and Sall4 were immunostained and the number of the triple positive cells were counted (n = 6 each). (B) Percentages for *Prx*GFP/Lhx2/Sall4-triple positive cells in cultures at Day8. (C) Other LP markers, Nmyc, Tfap2c and Msx1/2, were also stained. Error bars represent SD. Scale bar, 100 μm in (A).

**Supplemental figure 4**

**Transcriptomic comparison of the early limb bud to neighboring lateral plate mesodermal tissue, Related to** **Figure 1**

(A) Principal component analysis of five transcriptomic data sets. FL and HL bud expression values cluster closely together. Separation of other three data sets occurs across principal component 1 (PC1) and principal component 2 (PC2). (B) Top five most statistically significant enriched gene ontology classifications for top 100 genes associated with PC1 and PC2.

**Supplemental figure 5**

**Whole-mount views of HA-gels with 17 factors overexpressing cells (18-1 dropout assay), Related to** **Figure 2**

Each factor was withdrawn from the pools one by one to see which factor was critical for *Prx*GFP induction. Scale bar, 2 mm.

**Supplemental figure 6**

**Lin28a is a key factor for induction of limb marker genes, Related to** **Figure 2**

**(A)** 7 (Hoxd10, Tfap2a, Lhx2, Etv4, Prdm16, Zbtb16, and Lin28a) -1 factor dropout assay showed that every factor from the pools was critical for *Prx1*-GFP induction. When Lin28a was removed, the GFP score was the lowest and Sall4 proteins were not detected. (B) 7-2 dropout assay was performed. Note that GFP scores were decreased when Lin28a and one additional factor were withdrawn from the pools. (C) Single factor assay, in which only one factor was used to infect NonLFs, was conducted. Lin28a yielded the highest GFP score, and induced expression of Sall4, bsut not Lhx2. Error bars represent SD. Scale bars, 100 μm in (A), (C).

**Supplemental figure 7**

**Size reduction occurs after the reprogramming, Related to** **Figure 2**

(A) FACS profiles of NonLFs, PZL-reprogrammed cells and LPs from E9.5 *Prx1*-GFP embryos. FSC-A indicates surface area of cells. (B) Area of DAPI signals of Control (no virus condition), PZL-reprogrammed cells, or 3D cultured limb progenitors was measured (n = 50 cells each). \*\*\*\**p* < 0.0001, one-way ANOVA. Error bars represent SD. Scale bar, 100 μm in (B).

**Supplemental figure 8**

**Expression analyses of *Egr1* mRNA in the chicken embryos and Egr1 proteins in the mouse forelimb, Related to** **Figure 3**

(A) mRNA expression patterns of *Sall4*, *Lin28a*, *Lin41* and *Egr1* at the FL forming region of HH15 and FL buds of HH19 chicken embryos. *Egr1* was not present at HH15 (a black arrowhead) while it was detected at HH19 (a yellow arrowhead). (B) Egr1 (green) and MHC (red) were visualized in E13.5 mouse FL. Egr1 signals were localized at the end of myofibers as marked by yellow arrowheads. Scale bar, 500 μm in (B).

**Supplemental figure 9**

**Expression analysis for *Prx1*-GFP/tdTomato in the reprogrammed cells, Related to Figure 4 and 7**

Schematic representation of the strategy to induce *Prx1*-GFP and tdTomato by reprogramming. GFP and tdTomato expression were investigated in a cross section of an E9.5 *Prx1*-GFP/tdTomato reporter embryo, NonLFs, Control cells, PZL- and PZLL-reprogrammed cells. Control cells were infected with viruses carrying no transgene. Control, PZL- and PZLL-reprogrammed cells were cultured for 14 days as depicted in Fig. 4A. Scale bar, 100 μm.

**Supplemental figure 10**

**Representative FACS profiles of samples used for scRNA-Seq analyses, Related to** **Figure 5**

(A) Mouse cells transfected with PZL, cultured in CFRSY/HA condition. (B) Mouse cells transfected with no transgene (empty viruses), cultured in CFRSY/HA condition. (C) Fresh E9.5 LPs from *Prx1*-GFP mice. (D) mouse cells transfected with PZLL, cultured in CFRSY/Matrigel condition. (E) Mouse cells transfected with PZL, cultured in CFRSY/Matrigel condition. (F) Mouse cells transfected with no transgene (empty viruses), cultured in CFRSY/Matrigel condition. (G) E9.5 primary mouse cells cultured in CFRSY/Matrigel condition. (H) Human cells transfected with PZLL, cultured in CFRSY/Matrigel condition. (I) Human cells transfected with PZL, cultured in CFRSY/Matrigel condition. (J) Human cells transfected with no transgene (empty viruses), cultured in CFRSY/Matrigel condition. DAPI and DRAQ5 were used to mark dead and vital cells, respectively. The *Prx*GFP+ cells were sorted as reprogrammed cells. (m): mouse cells, (h): human cells. PZL refers to Prdm16+Ztbt16+Lin28a (3-factor lentiviral expression). PZLL refers to Prdm16+Ztbt16+Lin28a+Lin41(Trim71) (4-factor lentiviral expression).

**Supplemental figure 11**

**Aggregate cell-level statistics, Related to** **Figure 5**

(A) Cell counts per library, (B) UMI counts per cells, (C) Genes per cell, (D) mitochondrial fraction (UMI counts of mitochondrial genes divided by the total UMI counts), (E) ribosomal gene fraction (UMI counts of ribosomal genes divided by the total UMI counts). 10X Genomics v3 and InDrop technology have very different RNA capture rate, thus UMI counts as well as gene coverage. Therefore, each statistic was separated into the two technological batches. Matrigel-related samples (3D) were processed with 10X Genomics v3 technology, whereas the hyaluronan (HA)-related reprogramming samples were processed with InDrop technology. Red dots represent the median values for each sample annotation. PZL refers to Prdm16+Ztbt16+Lin28a (3-factor lentiviral expression). PZLL refers to Prdm16+Ztbt16+Lin28a+Lin41(Trim71) (4-factor lentiviral expression).

**Supplemental figure 12, Related to** **Figure 6**

**Expression of the reprogramming genes and *PrxGFP* in the UMAP**

(A) Four panels highlight cells from specified sample sources, with color contours representing the density of the corresponding sample source.

(B) Expression of PZLL genes, select limb patterning genes and fraction of transgene expression in UMAP coordinates. For specific genes, the values are log-transformed, UMI counts normalized by the total UMI counts of a cell. Fraction of transgenes represent the total number of UMIs attributed to the potential transgenes (includes EGFP, Woodchuck Hepatitis Virus Posttranscriptional Response element (WPRE) counts as well as human Lin41 (hLin41) UMI counts, with the addition of Prdm16, Ztbt16, Lin28a UMI counts, where the endogenous to transgene cannot be distinguished) to total UMI counts for a given cell and the maximum is 1. Maximum value for a given coordinate.

(C) Dot plot of patterning genes in all clusters. All expression level in natural-log transformed UMI counts normalized by the total UMI counts per cell, maximal expression. PZL refers to Prdm16+Ztbt16+Lin28a (3-factor lentiviral expression). PZLL refers to Prdm16+Ztbt16+Lin28a+Lin41(Trim71) (4-factor lentiviral expression).

**Supplemental figure 13, Related to Figure 5**

**The effect of 3D culture condition by comparing LPCs in 3D cultured condition for 8 days to LPCs harvested directly from corresponding stages**

(A) Left panel: Volcano plot comparing cultured limb progenitors (LPCs) and primary LPCs from rLPC/LPC (3D), LPC (E9), and LPC (E10) clusters (Leiden, resolution=0.2). Adjustment of p-values were performed by pseudobulk aggregation of expression data by independent E9.5-E10.5 samples that comprise more than 100 cells for the cluster 1, 4, 7 using Benjamini-Hochberg adjustment (3D cultured condition: n=2, Immediately harvested: n=7). Red dotted line is threshold of adjusted p-value=0.1. All genes more than 20 log fold changes, likely due to zero counts in one contrast, are put into infinity for better visualization. Right panel: UMAP plot showing the cluster and cells used for differentially expressed gene analysis in (A). Left panel: Bar plot of geneset enrichment analysis of differentially expressed gene lists from (A). k/K is the fraction of genes of a given gene set overlapping with the differentially expressed gene lists (cut-off adjusted p-value = 0.1). Right panel: -log10 of FDR q-value for the overlap. Select gene sets from MsigDB (Subramanian et al. 2005; Liberzon et al. 2011). (C) Gene Ontology (GO) term enrichment analysis of differentially expressed gene lists from (A), with Top 20 GO terms arranged by FDR q-value.

**Supplemental figure 14, Related to** **Figure 6**

**High-resolution clustering of scRNA-seq cells for Waddington Optimal Transport (WOT) analysis**

(A) UMAP plots of infected and 3D cultured cells used for scRNA-seq analysis split by sample date and the type of infection. All primary cells were excluded. (B) Cell cycle, Apoptosis gene set z-scores calculated for Waddington Optimal Transport (WOT) analysis with other Gene Ontology (GO) term gene sets and independently calculated G2M/S Scores and ribosomal fractions for reference. GO_FL_MORPHO (Embryonic forelimb morphogenesis, GO: 0035115), GO_JOINT_DEVO (Embryonic skeletal joint development, GO: 0072498), GO_TENDEON_DEVO (Tendon development, GO: 0035989), GO_CHONDRO_DEVO (Chondrocyte development, GO:0002063). (C) Left panel: UMAP plot with cells colored by high-resolution leiden cluster annotation (resolution=0.4) with circled labels positioned at the center of corresponding clusters. Right panel: Violin plots of select markers for the high-resolution leiden clusters (resolution=0.4). Only the expression levels of infected cells are shown. Bottom panel: Violin plot of expression of *Osr1* and *Acta2*, showing an overlap of a small *Acta2+ Osr1+* primary cells from E12.5 overlapping with the r2 cluster. (D) Alluvial diagrams showing the inferred transition and growth/contraction of infected cells in 3D culture from NonLFs to the Day 14 highlighted by the color of intermediate and final fate of the four rLPC sub-clusters. PZL refers to Prdm16+Ztbt16+Lin28a (3-factor lentiviral expression). PZLL refers to Prdm16+Ztbt16+Lin28a+Lin41(Trim71) (4-factor lentiviral expression).

**Supplemental figure 15, Related to** **Figure 6**

**High-resolution clustering of scRNA-seq cells for Waddington Optimal Transport (WOT) analysis**

(A) Changes of mean expression levels of individual genes at a given time point weighted by the probability of the final fate (rLPC or Transit) inferred by WOT. All expression level in natural-log transformed UMI counts normalized by the total UMI counts per cell. rLPC refers to reprogrammed limb progenitors. Transit refers to all cells with the cluster annotation of (A1, A2, T1, T2, T3).

(B) Changes of mean expression levels of individual genes at a given time point weighted by the probability of the final fate for individual rLPC fates (r1, r2, r3, E9). All expression level in natural-log transformed UMI counts normalized by the total UMI counts per cell. PZL refers to Prdm16+Ztbt16+Lin28a (3-factor lentiviral expression). PZLL refers to Prdm16+Ztbt16+Lin28a+Lin41(Trim71) (4-factor lentiviral expression). WPRE refers to Woodchuck Hepatitis Virus Posttranscriptional Response element, representing lentiviral expression level.

**Supplemental figure 16**

**Overexpression of PZL induces expression of LP marker genes in human adult fibroblasts**

(A) Control (no transgene) virus- or PZL-infected human dermal fibroblasts (HDF) were cultured on Matrigel in the presence of CFRSY for 18 days. (B, C) HDF, Control and PZL-overexpressing cells were stained with SALL4 and LHX2 antibodies (B), or NMYC and EGR1 antibodies (C). (D) After the PZL-expressing cells were cultured for 18 days, the cells were grafted into HH20 chicken FL buds, and then the grafted limbs were harvested at HH32, 4 days after the manipulation. A few grafted PZL cells were integrated in cartilage and became Sox9 positive (a yellow arrowhead), whereas control HDF do not differentiate into chondrocytes (n = 3 limbs each). Scale bars, 100 μm in (B), (C), (D). PZL refers to Prdm16+Ztbt16+Lin28a (3-factor lentiviral expression). PZLL refers to Prdm16+Ztbt16+Lin28a+Lin41(Trim71) (4-factor lentiviral expression).

**Supplemental figure 17**

**scRNA-Seq characterization of the human cells reprogrammed by PZL or PZLL**

(A) UMAP plot of single cell transcriptome embedding of human cells infected with empty, PZL-, PZLL-reprogramming factors and human dermal fibroblast (HDF) control.

(B) Violin plots of major markers for NonLFs, LPs in Matrigel cultured human cells with empty, PZL-, PZLL-reprogramming factors. (C) Combined UMAP embedding of mouse and human single cell transcriptomes. The four panels highlight cells from specified sample sources, with color contours representing the density of the corresponding sample source. PZL refers to Prdm16+Ztbt16+Lin28a (3-factor lentiviral expression). PZLL refers to Prdm16+Ztbt16+Lin28a+Lin41(Trim71) (4-factor lentiviral expression).

**Supplemental Table 1**

**List of all scRNA-Seq libraries, Related to** **Figure 5**

All inDrops and 10X v3 libraries used in the manuscript. The lower mapping rate for the inDrop libraries stems from insufficient cleaning up of short primer-dimers in the library. The numbers under quality control (QC) process represent the number of cellular barcodes for each library. “Initial CB” column represent the number of cellular barcodes (CB) suggested by the 10X cellranger/dropEst pipeline. “After QC” column represent the remaining cellular barcode after cut-off of primarily mitochondrial content and gene count per cell. “Relevant Cell type” column represent the remaining cellular barcodes, after clustering and marker analysis for each library and removing irrelevant, contaminating cell types, such as immune cells, muscle cells. “Singlet” column represent the number of remaining cellular barcodes after putative doublets were removed via Scrublet algorithm (Wolock et al. 2019). Prefix D means day after infection and culture, prefix E means mouse embryonic time point post coitum, PZL refers to Prdm16+Ztbt16+Lin28a (3-factor lentiviral expression). PZLL refers to Prdm16+Ztbt16+Lin28a+Lin41(Trim71) (4-factor lentiviral expression). HA refers to hyaluronan-based culture, F0 refers to Empty lentiviral infection control. NonLFs refers to non-limb fibroblasts.

**Supplemental Table 2**

**Differentially expressed genes for clusters in resolution=0.2 and resolution=0.4**

(A) Differentially expressed genes contrasting each broad cluster (leiden cluster resolution=0.2) to NonLF cluster, results related to Fig. 5B

(A) (B) Differentially expressed genes contrasting cells from reprogrammed limb progenitors (rLPCs) to 3D cultured limb progenitors (LPC), related to Fig. 5C.

(B) Differentially expressed genes contrasting cells from primary limb progenitor origin in different culture condition (3D cultured primary vs Immediately harvested primary), related to Fig. S13A

(C) Differentially expressed genes contrasting each fine cluster (leiden cluster resolution=0.4) to NonLF cluster, adjusted p-value cut-off of 0.1, results related to Fig. S14C.

Column specifications:

- name1/name2 : The source that is compared to each other. There are three distinct comparisons: PZL vs primary, PZLL vs primary, PZLL vs PZL. PZL refers to Prdm16+Ztbt16+Lin28a (3-factor lentiviral expression). PZLL refers to Prdm16+Ztbt16+Lin28a+Lin41(Trim71) (4-factor lentiviral expression). Primary refers to primary limb progenitors cultured in 3D culture condition.
- feature : Gene symbol
- pval : the p-value of the quasi-likelihood ratio test
- adj_pval : the adjusted p-values based on the pseudo bulk procedure treating each captured library as distinct source, not the individual cells
- f_statistic : the F-statistics
- df1 : the degrees of freedom of the test
- df2 : the degrees of freedom of the fit
- lfc : the log2-fold change.
- For more specifics, refer to glmGamPoi package test_de function.

**Supplemental table 3**

**Differentially expressed genes for trajectories**

(A) Weighted t-test results from PZL-infected as well as PZLL-infected cell trajectories comparing successfully reprogrammed limb progenitor fate (rLPC) to transit fate (Transit). Related to Fig. 6D, E, Fig. 15A

(B) Weighted t-test results from PZL-infected as well as PZLL-infected cell trajectories comparing each r2 limb progenitor fate to the rLPC states closer to earlier limb progenitors (r3/E9). Related to Fig. 15B

Column specifications:

- All suffixes refer to the statistics for a particular dataset. PZL refers to Prdm16+Ztbt16+Lin28a (3-factor lentiviral expression). PZLL refers to Prdm16+Ztbt16+Lin28a+Lin41(Trim71) (4-factor lentiviral expression).
- Day : The time point after infection and 3D culture which cells are selected to compare the trajectories.
- name1/name2 : The clusters that are compared to. rLPC (r1/r2/r3/E9 clusters aggregated), Transit (A1/A2/T1/T2/T3 clusters aggregated).
- feature : Gene symbol
- isTF : Whether the gene is a transcription factor, according to online resource of AnimalTFDB (Zhang et al., 2012)
- fold_change : Fold change between the two trajectories for the particular dataset
- mean1/mean2 : weighted mean expression value based on the fate probabilities of all cells.
- fraction_expressed1/fraction_expressed2 : weighted mean of the occurrence of the particular gene in the group
- t_score : weighted t-test score
- t_pval : weighted t-test p-value
- t_fdr : False Disvery Rate adjusting for the number of cells between groups

## STAR Methods

Detailed methods are provided in the online version of this paper and include the following:

- KEY RESOURCES TABLE
- LEAD CONTACT AND MATERIALS AVAILABILITY
- EXPERIMENTAL MODEL AND SUBJECT DETAILS

○ Mouse and chicken embryos
- METHOD DETAILS

○ Embryonic fibroblast isolation
○ Matrigel coating
○ Harvest and culture of limb progenitors
○ Quantitative PCR (qPCR)
○ Plasmid construction
○ Viral production
○ Reprogramming assay 1: Reprogramming for mouse embryonic non-limb fibroblasts using HA-hydrogels
○ Reprogramming assay 2: Reprogramming for mouse embryonic non-limb fibroblasts using Matrigel
○ Reprogramming assay 3: Reprogramming for human adult dermal fibroblasts using Matrigel
○ Immunostaining
○ Micromass culture and Alcian blue staining
○ Probes and *in situ* hybridization
○ *In ovo* electroporation
○ Tamoxifen and 4-Hydroxy Tamoxifen (4-OHT) treatment
○ Cell transplantation to chicken embryos
○ RNA-Seq library preparation
○ Dissociation and FAC-sorting of 3D cultured cells for single-cell RNA-Seq (scRNA-seq)
○ scRNA-seq library preparation: InDrops scRNA-seq
○ scRNA-seq library preparation: 10xGenomics scRNA-seq
- QUANTIFICATION AND STATISTICAL ANALYSIS

○ RNA-Seq analyses
○ scRNA-seq analyses
- DATA AND CODE AVAILABILITY

## Supplemental Information

Supplemental information can be found online at xxxxxxxxx.

## STAR★METHODS

### KEY RESOURCES TABLE

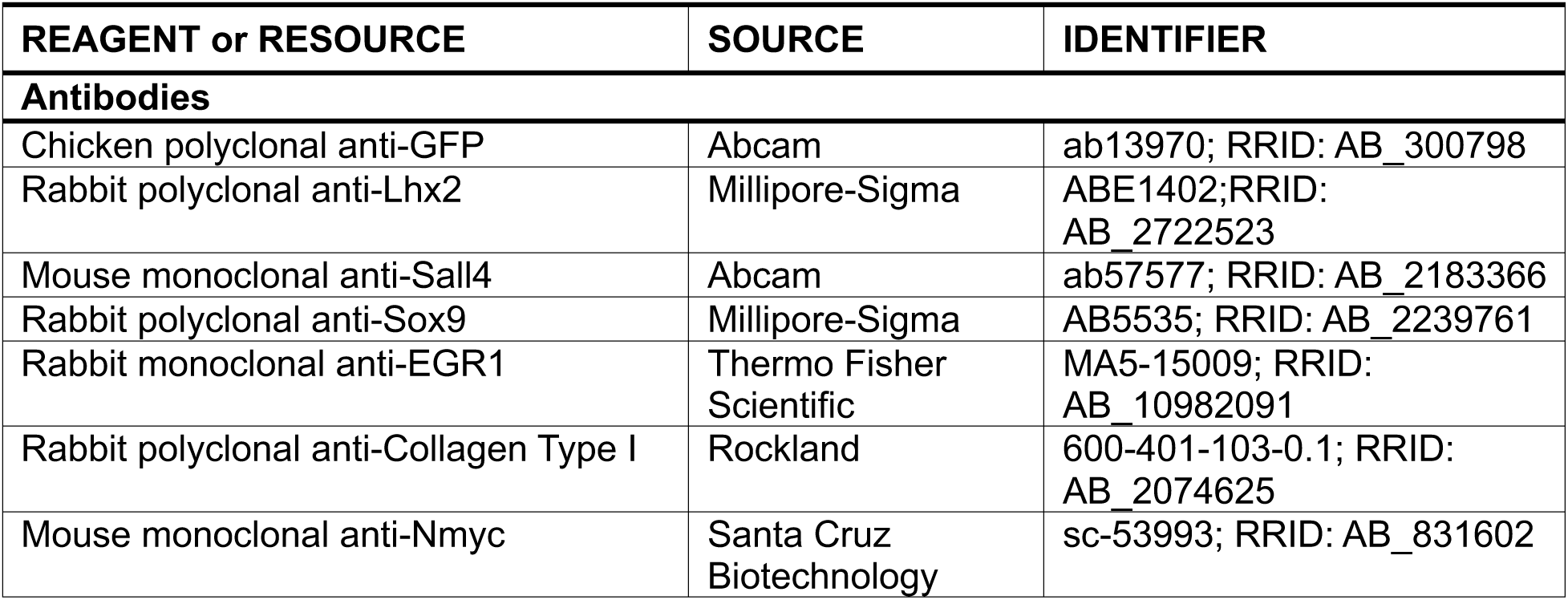
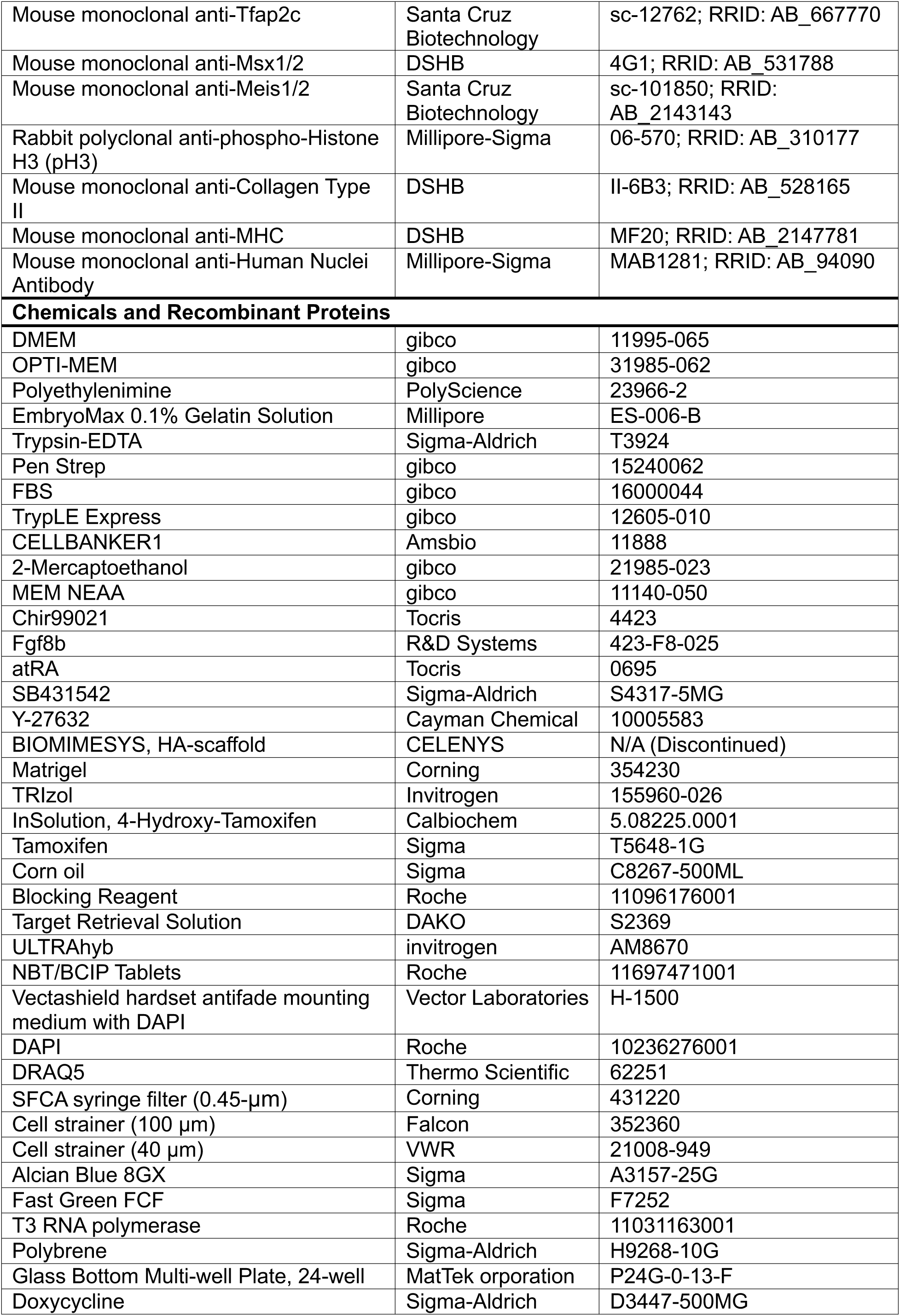
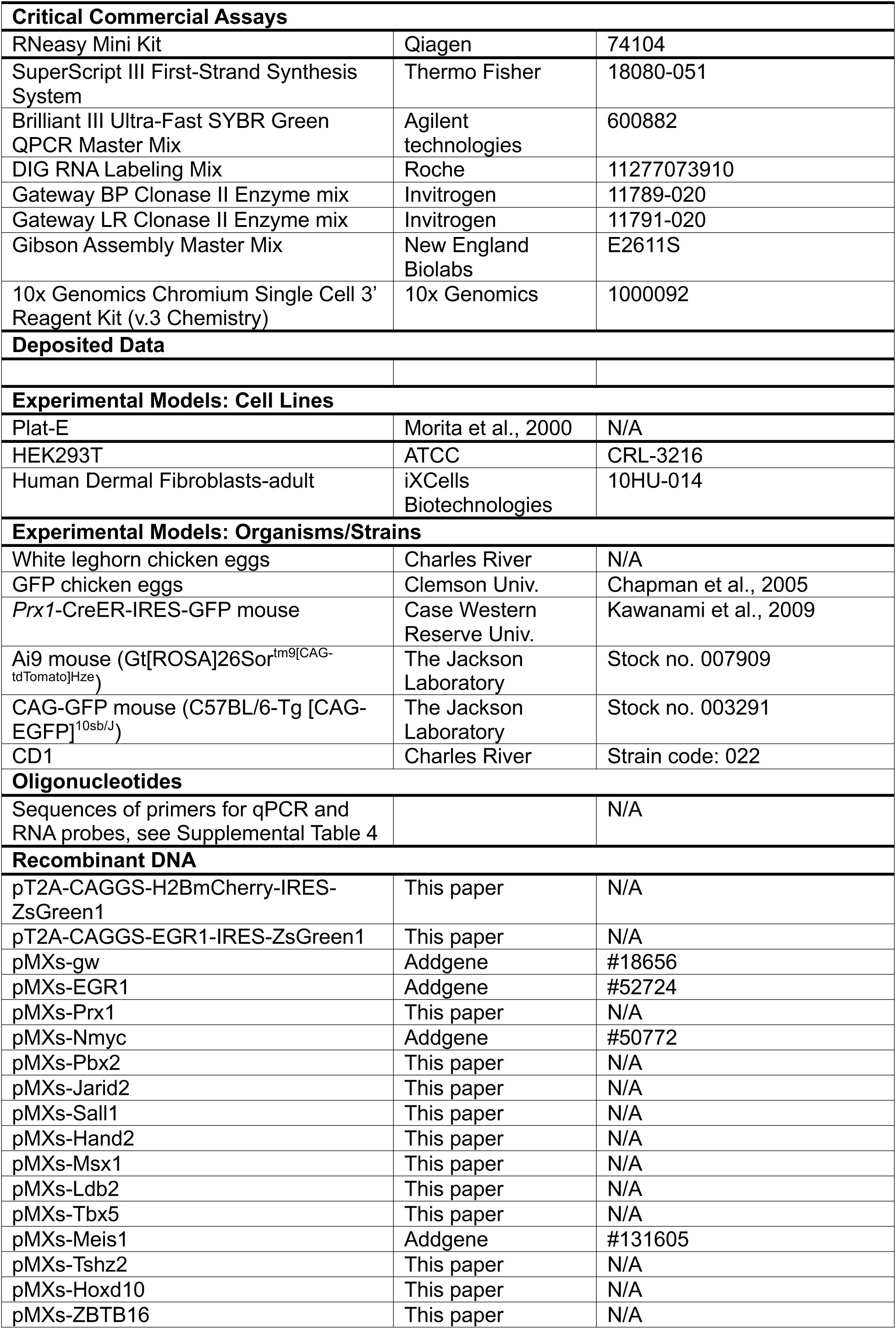
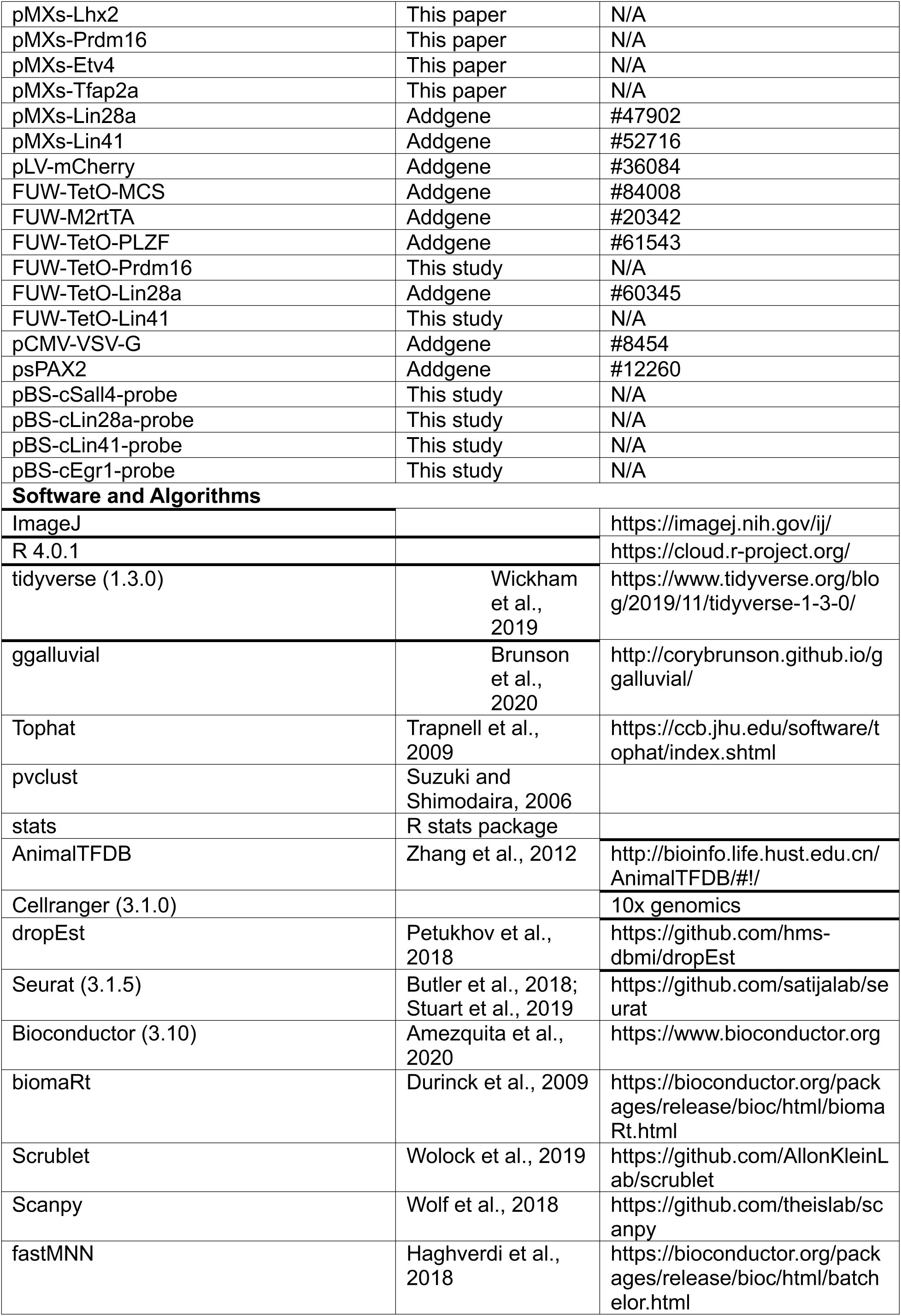
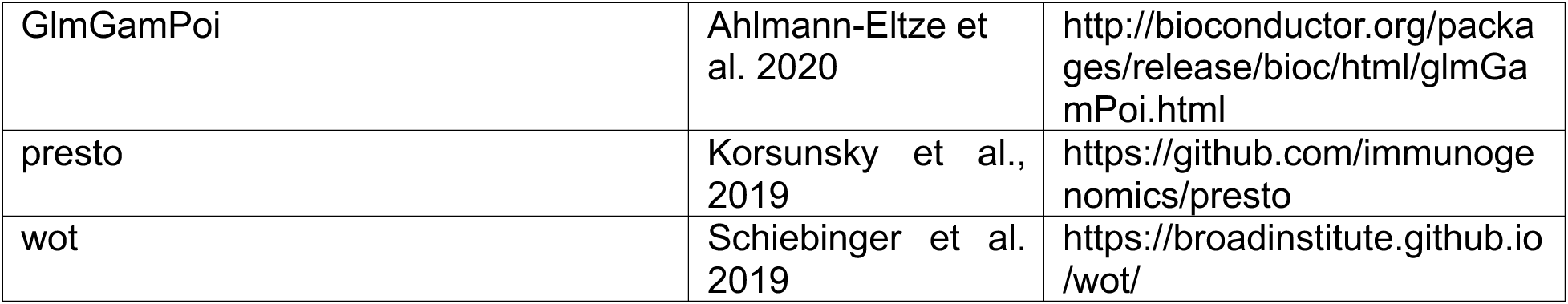

### LEAD CONTACT AND MATERIALS AVAILABILITY

Further information and requests for resources such as recombinant DNA plasmids generated in this study should be directed to and will be fulfilled by the Lead Contact, Clifford J. Tabin.

### EXPERIMENTAL MODEL AND SUBJECT DETAILS

#### Mouse and chicken embryos

Mouse colonies were maintained in the vivarium at the New Research Building of Harvard Medical School. *Prx1*-CreER-IRES-GFP (hereafter *Prx1*-GFP) mice were provided by Shunichi Murakami (Case Western Reserve University)(Kawanami et al., 2009). Ai9 (*Gt[ROSA]26Sor^tm9[CAG-tdTomato]Hze^*) and CAG-GFP (*C57BL/6-Tg [CAG-EGFP]^10sb/J^*) mouse strains were purchased from the Jackson Laboratory. Ai9 mice were crossed with *Prx1*-GFP mice to obtain *Prx1*-CreER-IRES-GFP:Rosa-CAG-LSL-tdTomato reporter embryos (*Prx1*-tdTomato). White leghorn eggs were obtained from Charles River. Chicken embryos were staged according to the Hamburger and Hamilton stages (HH) (Hamburger and Hamilton, 1951). All animal experiments were performed under the guidelines of the Harvard Medical School Institutional Animal Care and Use Committee.

### METHOD DETAILS

#### Embryonic fibroblast isolation

Embryonic fibroblasts were derived from E13.5 *Prx1*-GFP or *Prx1*-tdTomato embryos (the head, neck, limbs, lateral plate mesoderm derived tissues, and internal organs were discarded). The dissected embryos were minced with a razor blade and incubated in 0.25% Trypsin (Sigma) for 15 min. The suspension was plated in Gelatin (Millipore)-coated 15-cm tissue culture dishes in DMEM/10%FBS/1%Pen-Strep media (DMEM/FBS). The cells were grown at 37℃ in 5% CO_2_ until confluent, and GFP- or GFP/tdTomato-negative fibroblasts were collected by a FAC sorter Astrios (Beckman Coulter). After the sorted cells were grown until confluent, the cells were split once before being frozen (Passage 3).

#### Matrigel coating

200 ul of Matrigel (Corning) is diluted with 200 ul of chilled OPTI-MEM (gibco) (1:1 dilution), and the diluted Matrigel was placed in a well of a 24-well plate (Corning). The plate was incubated to be gelatinized in a cell culture chamber at 37℃ for 30 min.

#### Harvest and culture of limb progenitors

Forelimb (FL) buds from E9.5 *Prx1*-GFP mouse embryos or HH18 GFP-chicken embryos were dissected out and incubated in 0.25% Trypsin for 5-10 min at room temperature to loosen ectodermal tissues. After the surface ectoderm was removed by fine forceps, limb progenitors (LPs) were dissociated gently by pipetting and pelleted by centrifugation. The cells were re-dissociated by culture media, and LPs obtained from two limb buds were placed in one well of 24-well plate dishes, a hyaluronan (HA)-based hydrogel (CELENYS) or a well of Matrigel-coated 24-well plate dishes. To make the LP culture media (CFRSY media), DMEM/FBS was supplemented with 3 μM Chir99021 (Tocris), 150 ng/ml Fgf8 (R&D Systems), 25 nM Retinoic acid (Tocris), 5 μM SB431542 (Sigma-Aldrich), 10 μM Y-27632 (Cayman Chemical), 55 μM 2-Mercaptoethanol (gibco), and MEM Non-Essential Amino Acids Solution (100X, NEAA, gibco). The media was changed every other day until Day6, and then changed every day until Day10.

#### Quantitative PCR (qPCR)

RNA was extracted using Tryzol (Invitrogen) or RNeasy Mini kit (Qiagen). For qPCR of *Lin28a*, RNAs were extracted from FL, HL and flank mesenchyme located between FL and HL buds at E9.5 CD1 mouse embryos by using RNeasy Mini kit. To recover RNA from the cells cultured in the HA-hydrogels, the cells in the gels were lysed in 1 ml Trizol (for 1 to 5 hydrogels) by vortexing for 5 min. 200 μl of Chloroform (Sigma-Aldrich) was added and vortexed for 10 sec, and then incubated for 3 min at room temperature. After centrifugation (10,000 g, 20 min, 4℃), aqueous phase was collected, and 500 μl isopropanol was added. After centrifugation and two washes with 80% ethanol, RNA pellets were dissolved in RNase-free water and kept at -80℃ until use. The collected RNA was reverse-transcribed by SuperScript III First-Strand Synthesis System (Thermo Fisher). PCR reaction was performed by using Brilliant III Ultra-Fast SYBR Green QPCR kit (Agilent) and CFX Touch Real-Time PCR Detection System (Bio-Rad). Relative expression levels were calculated by the ΔΔCq method. Sequences (5’-3’) of primers for qPCR are described in Table S4.

#### Plasmid construction

The coding regions of candidate genes were PCR-amplified from mouse embryo derived cDNA or purchased clones (Thermo Scientific). The PCR-amplified sequences were cloned into pDONR221 using the Gateway BP reaction mix (Invitrogen). The resulting entry clones were then recombined with pMXs-gw (Gift from Shinya Yamanaka; Addgene #18656) using the Gateway LR reaction mix (Invitrogen). For FUW-TetO-Prdm16 and FUW-TetO-Lin41, cDNAs of Prdm16 and Lin41 were amplified by PCR from pMXs-Prdm16 and pMXs-Lin41 inserted to FUW-TetO-MCS (Addgene #84008) using Gibson Assembly Mix (New England Biolabs), respectively. To obtain pT2A-CAGGS-H2B-mCherry-IRES-ZsGreen1 and pT2A-CAGGS-EGR1-IRES-ZsGreen1, cDNAs of H2B-mCherry and EGR1 were integrated into pT2A-CAGGS-IRES-ZsGreen1 (Atsuta and Takahashi, 2016). For pBS-cSall4, pBS-cLin28a, pBS-cLin41 and pBS-cEgr1, the sequences amplified by PCR from HH18 or HH24 FL cDNA libraries that were generated by SuperScript III First-Strand Synthesis System (Thermo Fischer) were cloned into pBS-D (a gift from Dr. Daisuke Saito [Kyushu University]).

#### Viral production

Plat-E cells (Morita et al., 2000) were grown to 60-70% confluency in 10-cm dishes. pMXs-based retroviral vectors were transfected using Polyethylenimine (PEI, PolyScience). 30 μl of PEI (1 mg/ml) was diluted in 70 μl OPTI-MEM and incubated for 5 min at room temperature. 10 μg plasmid DNA was added to 100 μl OPTI-MEM, and then PEI and plasmid DNA solutions were combined and vortexed vigorously. The mixture was incubated for 30 min, and was added to the Plat-E cells. The cells were incubated for 24 hrs, and the media was replaced with 5 ml of fresh DMEM/FBS. The cells were incubated for another 24 hrs. 48 hrs after the initial transfection, the supernatant was collected and filtered. For production of lentiviruses, 293T cells were cultured up to 50-60% confluency in 10-cm dishes. 40 μl of PEI was diluted in 60 μl OPTI-MEM and incubated for 5 min at room temperature. 7.5 μg plasmid DNA carrying the reprogramming factor, 4.5 μg psPAX2 and 1.5 μg VSV-G plasmids were added to PEI solution, and the transfectant was incubated for 30 min. Then, the mixture was added to 293T cells, and 48 hrs after the transfection, the supernatant was harvested and filtrated through 0.45-μm SFCA syringe filters (Corning).

#### Reprogramming assays

##### Reprogramming for mouse embryonic fibroblasts using HA-hydrogels

At 60-70% confluency, mouse embryonic fibroblasts (*Prx1*-GFP negative) were cultured in the supernatant of retroviruses carrying the candidate factors for 24 hrs in the presence of Polybrene (8 μg/ml; Sigma-Aldrich) at 37℃ (Day 0), and the media was replaced with DMEM/FBS containing 2-Mercaptoethanol and NEAA (Day 0). 48 hrs after viral infection, the media was supplemented with 3 μM Chir99021, 150 ng/ml Fgf8, 25 nM Retinoic acid, 10 μM Y-27632, 55 μM 2-Mercaptoethanol, and Non-Essential Amino Acids (CFRY media; Day2). 48 hrs after CFRY administration, the viral infected cells were detached by Trypsin/EDTA, and the cells from each well of 24-well plates were suspended in 20 μl of CFRSY media (CFRY plus 5 μM SB431542). Subsequently, the cell suspension was loaded on the top of the HA-gels, and the gels were incubated for 30 min at 37℃. After incubation, the HA-gels were placed in 200 μl of CFRSY media, and the media was changed with the fresh CFRSY media every two days from Day4 to 10, every day from Day11 to 14. See also the schematics in Fig. 1G.

##### Reprogramming for mouse embryonic fibroblasts using Matrigel

At 60-70% confluency, GFP/tdTomato-negative fibroblasts from *Prx1*-tdTomato mice were cultured in the supernatant of lentiviruses carrying Prdm16, Zbtb16, Lin28a, and Lin41 (PZLL) for 24 hrs in the presence of Polybrene (8 μg/ml) at 37℃ (Day -1). The media was replaced with DMEM/FBS containing 2 μg/ml of Doxycycline (Dox; Sigma-Aldrich), 2-Mercaptoethanol and NEAA (Day 0). Next day the media was replaced with CFRY media containing Dox (CFRYD media; Day 1). 48 hrs after Dox administration, the cells were dissociated with TryPLE Express, and plated on Matrigel. The media was supplemented with CFRSYD media (Day 3), and was changed with the fresh CFRSYD media every two days from Day4 to 10, every day from Day11 to 14. 4-hydroxy tamoxifen (Calbiochem) was added to the media at Day 12 and Day13, to induce *Prx1*-tdTomato. See also the schematics in Fig. 4A and S9.

##### Reprogramming for human adult fibroblasts using Matrigel

Similarly to mouse cell reprogramming, human fibroblasts (iXCells Biotechnologies) were transduced with lentiviruses to misexpress PZLL at 60-70% confluency. After 2day-culture of DMEM/FBS/Dox and another 2day-culture with CFRY/Dox, the cells were transferred onto Matrigel bed and cultured for additional 14 days with CFRSY/Dox media. The total culture term was 18 days.

#### Immunostaining

For immunohistochemical staining, the following antibodies were used as described previously (Atsuta et al., 2019): anti-GFP (1:1000; Sigma), anti-Lhx2 (1:500; Millipore-Sigma), anti-Sall4 (1:500; Abcam), anti-Sox9 (1:500; Millipore-Sigma), anti-EGR1 (1:250; Thermo Fisher), anti-Collagen type I (1:100; Rockland), anti-Nmyc (1:500; Santa Cruz Biotechnology), anti-Tfap2c (1:500; Santa Cruz Biotechnology), anti-Msx1/2 (1:100; DSHB), anti-pH3 (1:500; Millipore-Sigma), anti-Collagen type II (1:100; DSHB), anti-MHC (1:50; DSHB), and anti-Human nuclei (1:250; Millipore-Sigma). For staining of Col2A1, an antigen retrieval using Target Retrieval Solution (DAKO) was performed in advance of blocking. To stain the 3D-cultured cells embedded in the HA-gel or Matrigel, the cells in the gels were placed in 1% PFA/PBS overnight at 4℃. The next day, the gels with the cells were incubated in 0.5% Triton X-100 (Sigma-Aldrich)/PBS for 15 min at room temperature, and then in 1% Blocking Reagent (Roche)/TNT buffer for 1 hr at room temperature, followed by primary and secondary antibody incubations. The stained cells were placed on a glass-bottom dish (MatTek), and images were taken by the confocal microscope LSM710 (Carl Zeiss).

#### Micromass culture and Alcian blue staining

Micromass culture and alcian blue staining were performed as previously described (Atsuta et al., 2019). Fibroblasts and LPs from E9.5 *Prx1*-GFP mouse forelimb (FL) buds, and *Prx1*-GFP positive reprogrammed cells were used to generate micromass cultures. ∼5 x 10^4^ cells per 20 μl of DMEM/FBS were dropped into each well of 96-well. After being attached, the cells were cultured in the presence of CFRSY for 2 days, and then in DMEM/FBS for 8 days.

#### Probes and *in situ* hybridization

Whole mount *in situ* hybridization for HH15 and HH17 chicken embryos was performed as described in (Tonegawa et al., 1997). cDNA sequences for chicken *Sall4*, *Lin28a*, *Lin41* and *Egr1* are described in Supplemental Table 4. RNA probes were transcribed using DIG-RNA labeling Mix (Roche) and T3 RNA polymerase (Roche), and the probes were detected with NBT/BCIP solution (Roche).

#### *In ovo* electroporation

The *in ovo* electroporation was performed as previously described (Atsuta et al., 2019). Briefly, eggs were incubated for approximately 54 hrs at 38℃. DNA solution was prepared at 4 μg/μl, and injected into the coelomic cavity of HH14 embryos. Three electric pulses of 50 V, 2 ms, were given, followed by 7 pulses of 5 V, 10 ms, with 10-ms interval between pulses (Super Electroporator NEPA21-type II, NEPA GENE).

#### Tamoxifen and 4-Hydroxy Tamoxifen (4-OHT) treatment

Tamoxifen was dissolved in corn oil (Sigma-Aldrich), and 1 mg of tamoxifen was given to E8.5 *Prx1*-tdTomato pregnant dams by intraperitoneal injections; 2 μM of 4-OHT (Calbiochem) was used for reprogramming experiments to activate CreER proteins.

#### Cell transplantation to chicken embryos

For cell injection, LPs from E9.5 CAG-GFP mouse FL, fibroblasts infected with lentiviruses carrying mCherry, and *Prx1*-tdTomato positive reprogrammed cells were used. The LPs form 10 FL buds were dissociated in 100 μl of DMEM/FBS. The mCherry-expressing fibroblasts and the tdTomato-reprogrammed cells were retrieved from one well of 24-well plates using TryPLE Express, and after pelleted, the cells were dissociated with 50 μl of DMEM/FBS. The cell suspension was injected in FL buds of HH20 chicken embryos, and the embryos were harvested at HH32.

#### RNA-Seq library preparation

Fertilized chicken eggs were incubated at 38℃. FL/HL buds and flank/neck mesenchyme were dissected from HH18/19 embryos. Neck tissue was located in the mesenchyme directly above the FL bud. Loose ectodermal tissues were removed and remaining mesenchyme was placed in TRIzol (Invitrogen) for RNA extraction. RNA-Seq on chick RNA was carried out as previously described (Christodoulou et al., 2014). Libraries were constructed without RNA or cDNA fragmentation and did not include normalization. Uniform amplification was achieved with amplification cycling before the reaction reached saturation, as determined by qPCR. Following Hi-Seq (Illumina) sequencing, reads were aligned using Tophat (version 1.4.0) (Trapnell et al., 2009).

#### Dissociation and FAC-sorting of 3D cultured cells before scRNA-seq

For sorting reprogrammed *Prx1-*GFP or *Prx1-*tdTomato cells, a FACS sorter Astrios (Beckman Coulter) or On-chip Sort HSG (On-chip Biotechnologies) was used. After washing with PBS, the cells cultured in the HA-gels or on Matrigel were incubated in TryPLE Express (gibco) for 30 min at 37℃. The cell suspension was pipetted with cut P1000 pipette tips every 10 min, to completely dissociate the cell clusters. The suspension was filtrated by 100 μm Cell strainers (Falcon) and 40 μm Cell strainers (VWR), and cells were pelleted by centrifugation (400 x g for 5 min). The pellets were dissociated by DRAQ5/DAPI in 0.1% BSA/PBS and incubated for 5 min before the sorting. . DRAQ5-positive, DAPI-negative cells were sorted for cells on reprogramming at day 2, 4, 8. For HA-gel reprogrammed cells at day 14, additional gating on GFP channel derived GFP-positive and GFP-negative samples. For Matrigel-derived day 14 reprogrammed cells for PZL-as well as PZLL-factors, only GFP-positive cells were collected. DRAQ5-positive, DAPI-negative, Matrigel-derived day 8 cultured E9.5 and E10.5 limb progenitors were collected. The E9.5 cultured limb progenitors were subject to 4-OHT, such that large fraction were tdtomato-positive, but the cells were collected regardless of tdTomato-positivity. DRAQ5-positive, DAPI-negative, tdTomato-positive cells were sorted for the limb mesenchyme cells for E10.5, E11.5 as well as E12.5 cells. For E9.5 limb progenitors, samples were collected without tdTomato gating to maximize yield.

#### Single-cell RNA-Seq library preparation

##### InDrops scRNA-seq

LPs were obtained from E9.5 and E10.5 mouse FL buds. HA-gel derived reprogrammed cells (PZL-factor) and empty controls, as well as CFSRY cultured NonLFs were collected and processed individually. cDNA library preparation was performed by Single Cell Core (HMS).

##### 10xGenomics scRNA-seq

FAC-sorted LPs were obtained from E9.5 and E10.5 *Prx1*-tdTomato mouse FL buds. All libraries included about 10-15% of MEF cells to mitigate batch effect. cDNA library preparation was performed by using 10x Genomics Chromium Single Cell 3’ (v.3 Chemistry; 10x Genomics) gene-expression kit, according to manufacturer’s instructions. Gel beads in emulsion (GEM) formation was performed with a Chromium Controller (10x Genomics; Biopolymer Facility at HMS). cDNA library was prepared in house.

#### Single-cell RNA-Seq sequencing

InDrops libraries were sequenced with Illumina Nextseq 500 platform, using paired-end reads with the read length configuration recommended by InDrops (61bp for transcript, 14bp for barcode and UMI, 8bp i7 index for part of barcode, 8bp i5 index for sample index). 10x Genomics libraries were sequenced with Illumina Nextseq 500 platform as well as Novaseq 6000 platform. For Nextseq 500, recommended configuration by 10x Genomics (28bp for cell barcode 1 and UMI, 8bp i7 index for sample index, 98bp for transcript) we used. For Novaseq, 150bp paired-end sequencing with sample i7 index were used (compatible with the 10x Genomics cellranger count matrix mapping software).

### QUANTIFICATION AND STATISTICAL ANALYSIS

#### RNA-Seq analyses

Analysis on transcriptome gene expression was conducted in R. The pvclust package (Suzuki and Shimodaira, 2006) was used to perform principal component analysis. The AnimalTFDB (Zhang et al., 2012) online resource was used to select transcription factors from the chick and mouse genomes.

#### Single-cell RNA-Seq analyses

##### Transcriptome annotation

For mouse samples, Ensembl release 98 mm10 transcriptome was used as base transcriptome annotation, with pseudogenes filtered from the GTF file using cellranger mkgtf command. For retroviral infected hyaluronan samples, transgenes for human Lin28a, EGFP (for Prx1GFP transgene) was added to generate custom transcriptome annotation for quantification for reprogrammed cells. For lentiviral infected Matrigel samples, transgenes for EGFP (for Prx1GFP transgene), rtTA, and human Lin41 as well as PLZF, and 3’UTR sequences of WPRE were added to generate custom transcriptome annotation for quantification for reprogrammed cells. The limb progenitor cells were subject to the same transcriptome annotation (yielding zero counts for the transgenes). All four human samples were multiplexed with mouse and chick samples (Supplementary Table 1). The chick data was not presented in this manuscript. Thus, for species demultiplexing, Ensembl release 99 hg38 transcriptome, Ensembl release 98 Gallus gallus-6.0 transcriptome and the filtered Ensembl release 98 mm10 transcriptome was merged using the cellranger mkgtf command to generate human-mouse-chick transcriptome for initial mapping for demultiplexing. For human-specific mapping, the filtered hg38 transcriptome with transgenes for EGFP, rtTA, and mouse Prdm16 and mouse Lin28a were added. For the limb progenitor samples processed with 10X genomics, tdtomato, EGFP transgenes were added for mapping.

#### InDrop preprocessing

Sequencing results were demultiplexed by dropTag from dropEst package (Petukhov et al. 2018). The demultiplexed reads were aligned with STAR aligner (Dobin et al. 2013). The aligned reads were split into forward and reverse alignment, since InDrops is directional. The resulting forward and reverse alignment files were quantified using dropEst package including directional UMI correction option (Petukhov et al. 2018) with transcriptome annotation split into forward and reverse direction to avoid mapping of antisense reads.

#### 10x data processing

Sequencing results were demultiplexed by cellranger and aligned using cellranger count (internally by STAR aligner (Dobin et al. 2013)). For the four libraries that needed species demultiplexing, cellular barcodes that had less than 5% of UMI counts from other species were selected for subsequent mapping with the corresponding species transcriptome (see above).

#### Quality control and clustering

Cellular barcodes with high mitochondrial content (>15%), high hemoglobin gene count (>10%) and low gene counts (<1,200) were filtered out. All libraries were subject to doublet detection via Scrublet (Wolock et al. 2019). The overall findings were not sensitive to the identified doublets. Batch effect was assessed by simply merging the individual UMI count matrices for clustering, which revealed dominant batch effect by technology (InDrop vs 10X) and time (the last 10X batch was separated by several months due to the pandemic). Thus, Seurat v3 integration procedure (SCTransform based) was applied (Stuart, Butler et al. 2019) with 30 dimensions for the individual batches. Further, cell cycle effect, a fraction of mitochondrial genes was regressed out. Principal component analysis (PCA) was performed on the integrated, scaled features for dimensional reduction and Uniform Manifold Approximation and Projection (UMAP) (McInnes et al. 2018) was used primarily for the cellular embedding coordinates. Leiden algorithm was applied on the neighbor graph with 10 iterations (Seurat default) to derive cluster boundaries (Traag et al. 2019). For all steps of clustering, the number of principal components were determined by observing the “elbow” of variance explained by the principal components, however, robustness of the relationship was confirmed by changing the number of principal components and deriving essentially similar relationship. Thus, 20 principal components were used for downstream processing. Resolution parameter of 0.2 was used for gross subdivision of all cells into 7 clusters (Fig. 5), and a resolution of 0.4 was used for leiden clustering for differential expression analysis and trajectory inference for Partition-based graph abstraction (PAGA) (Wolf et al. 2019) (Fig. 6). For presenting the embedding of human cell results only, the UMAP plot was based on fastMNN batch correction (Haghverdi et al. 2018).

#### Differentially expressed gene analysis, Gene set overlap analysis

Differentially expressed gene analysis (Fig. 5C, S13, Supplementary Table 2) were conducted with the glmGamPoi package (Ahlmann-Eltze et al., 2020) modelling the the batch effect as an additive latent variable and p-values were adjusted as pseudobulk procedure treating each biological samples as one unit rather than cells, yielding adjusted p-values (Benjamini-Hochberg). For Fig. 5C, cells from specific clusters were subsetted and further filtered such that the pseudobulk samples will have at least more than 100 cells, and instead of cluster labels, the sample origin (primary, PZL, PZLL) were used as a model variable. Similarly, for Fig. S13A, cells from specific clusters were subsetted and filtered as well and culture condition (Immediately harvested or 3D cultured for 8 days) were assigned for the samples as modelling variable. For those differentially expressed genes, the list were compared to the curated gene sets (Fig. S13B), or Gene Ontology (GO) terms (Biological process) for overlap by chance using MSigDB (Subramanian et al. 2005; Liberzon et al. 2011). To derive differentially expressed genes for Fig. 6D and Supplementary Table 3, simple weighted t-test based differential expression analysis provided by the Waddington Optimal Transport (WOT) analysis package was used, with the full reservation that the p-values will be artificially low.

#### Waddington Optimal Transport (WOT) analysis

The Waddington Optimal Transport analysis estimates the growth rate based on the cell cycle as well as apoptosis gene scores, calculated by z-score normalization (Schiebinger et al. 2019). Combat batch correction (Johnson et. al 2007) provided by scanpy framework was applied to the log-normalized UMI expression level before deriving the z-scores. The resulting cell cycle score as well as apoptosis score was used to infer the initial cell growth estimates, and growth fraction estimation as well as transport maps for control virus-infected time series, PZL (Prdm16+Ztbt16+Lin28a; 3-factor lentiviral expression)-infected time series, PZLL (Prdm16+Ztbt16+Lin28a+Lin41; 4-factor lentiviral expression)-infected time series were calculated separately with the following parameters: epsilon=0.05, lambda1=1, lambda2=50, growth_iteration=3. The choice of parameters were not sensitive for the overall findings. Since the day 8 PZL scRNA-seq had very low coverage, transcriptomes from day 8 PZLL-infected cells that do not show expression of transgene human Lin41 were included for the inference of this intermediate stage inference. Based on the transport maps, the ancestor and descendant relationship was calculated resulting in transition matrices between time points. The resulting transition tables were used to construct the alluvial diagrams used in Fig. 6B abd Fig. 14D, with the ggalluvial package (Brunson et al. 2020). The cell sets for each high resolution leiden clusters (resolution=0.4) at Day 14 were defined as final fates, and the fate probability, weighted mean expression at different time points for individual genes was calculated for Fig. 6D and Fig. S14, S15 and Supplementary Table 3.

#### Human/Mouse scRNA-seq data processing

The four human UMI count matrices were merged first and only orthologous genes (1:1 matching) from the human transcriptome based on biomaRt (Durinck et al., 2009) were translated into mouse genes. The resulting matrix were integrated with the mouse libraries treating the human libraries as a separate batch (SCTransform-based Seurat integration). All subsequent clustering steps were identical to the mouse-only analysis.

### DATA AND CODE AVAILABILITY

The datasets and code utilized in this study are available at GEO: GSE XXXXXXXX and on GitHub at https://github.com/YYYYYYYYYYY.

