## Supplementary Figures for "Direct Reprogramming of Non-limb Fibroblasts to Cells with Properties of Limb Progenitors"

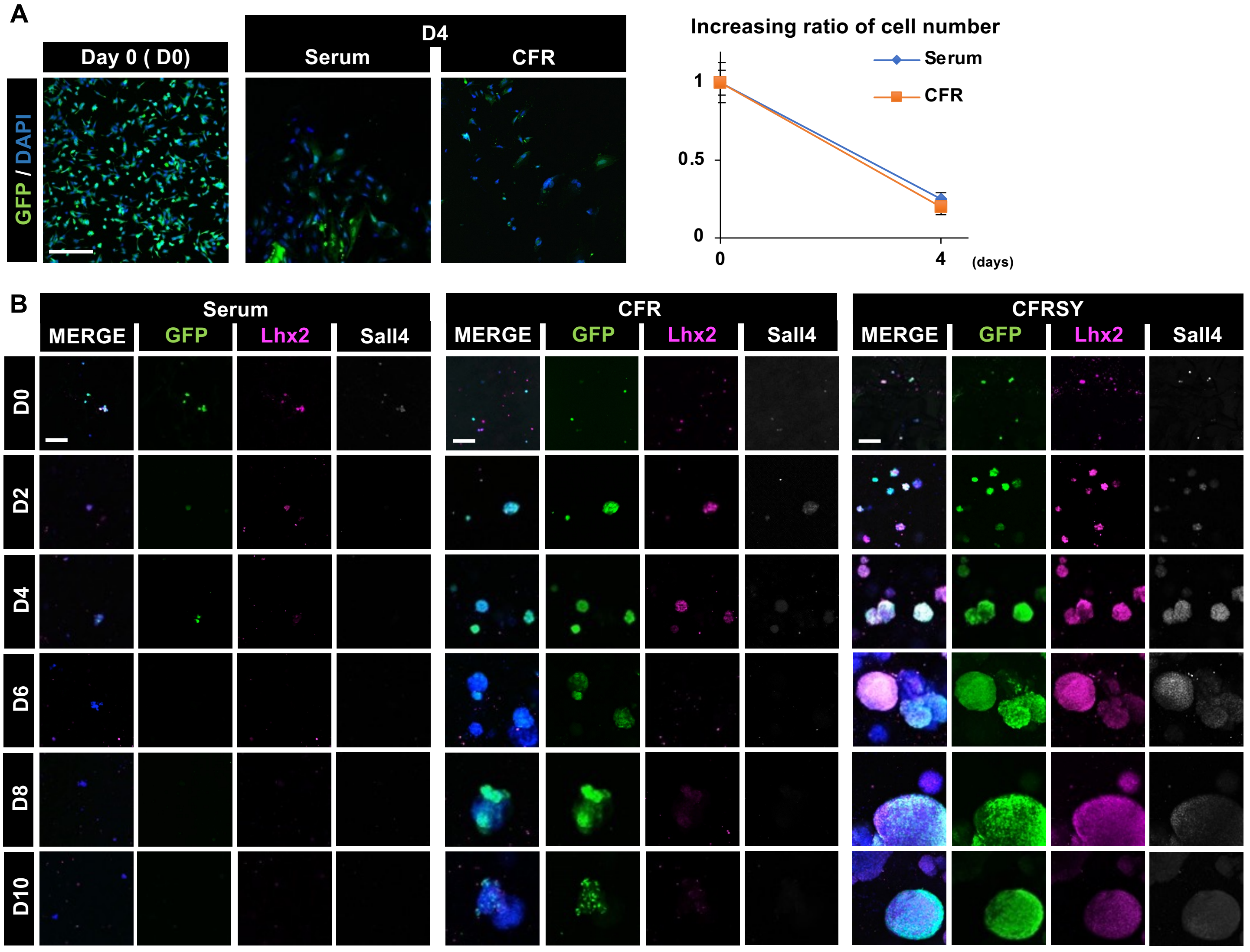


Fig. S1 Atsuta et al.

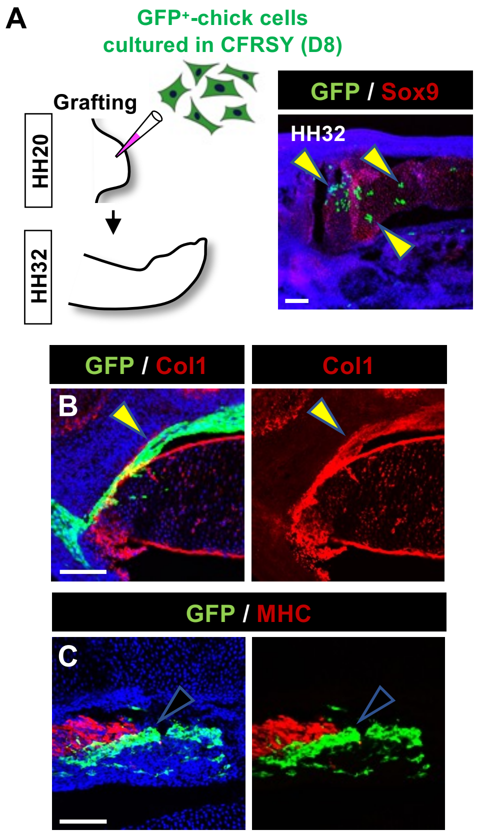


Fig. S2 Atsuta et al.

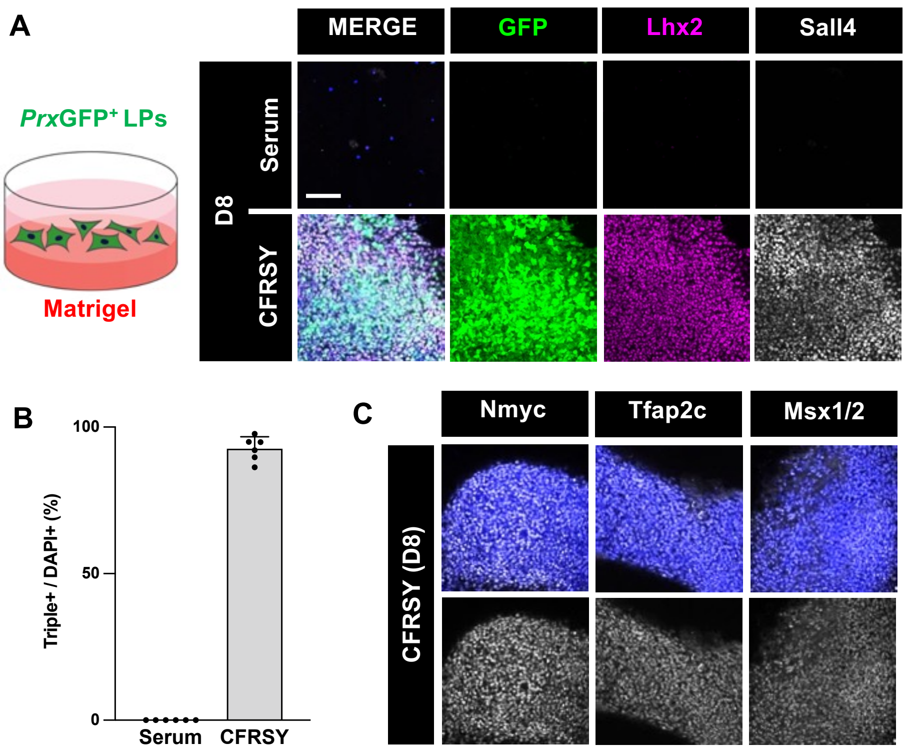


**Fig. S3 Atsuta et al.**

A

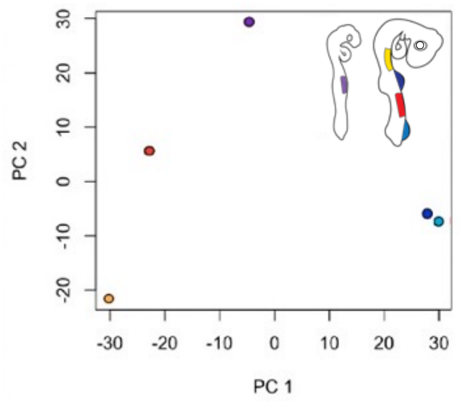


B

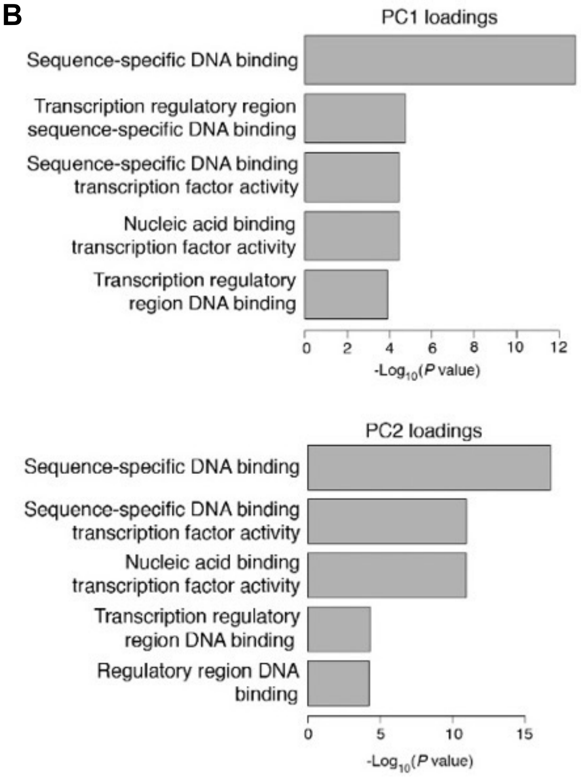


**Fig. S4 Atsuta et al.**

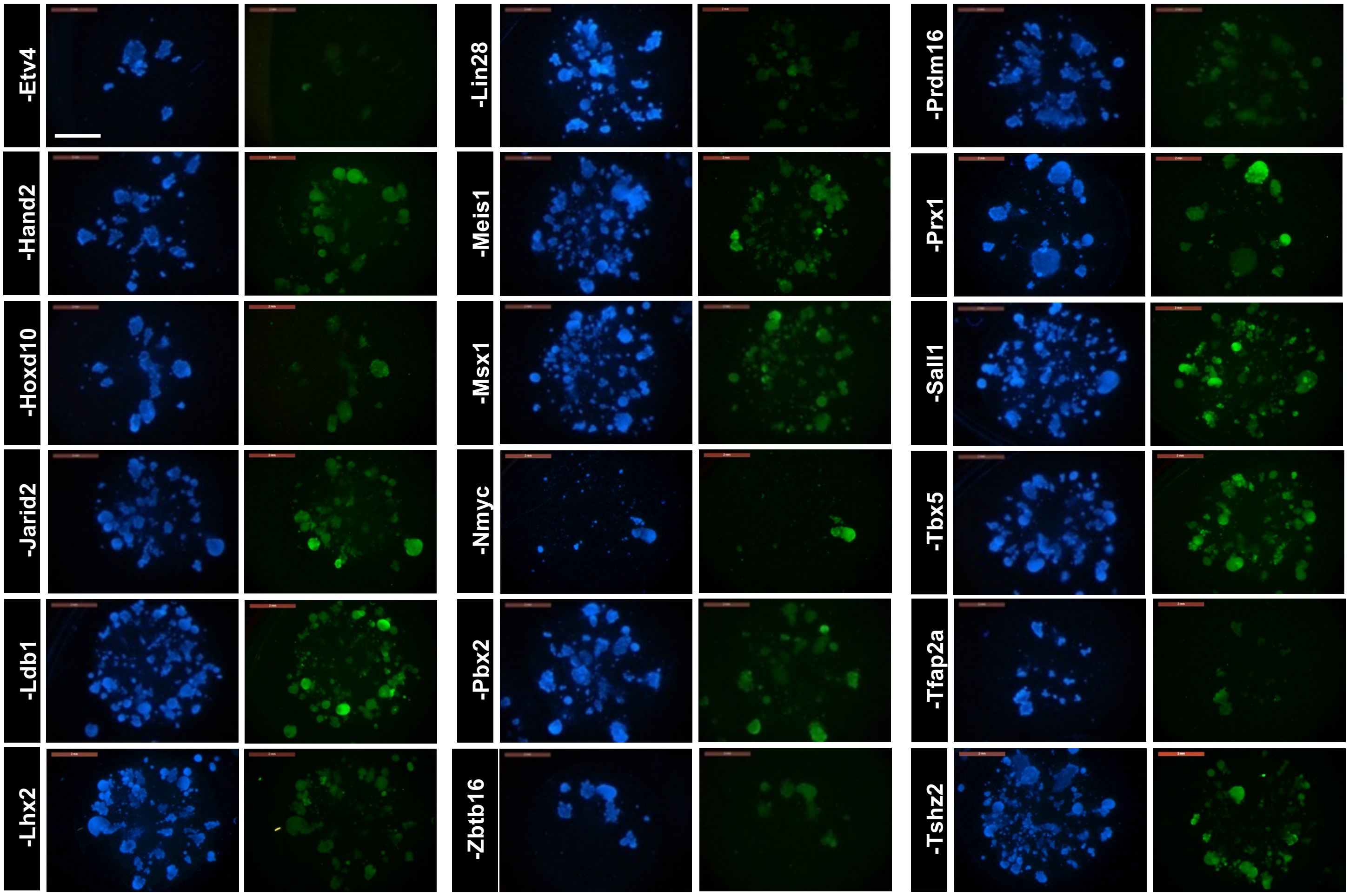


**Fig. S5 Atsuta et al.**

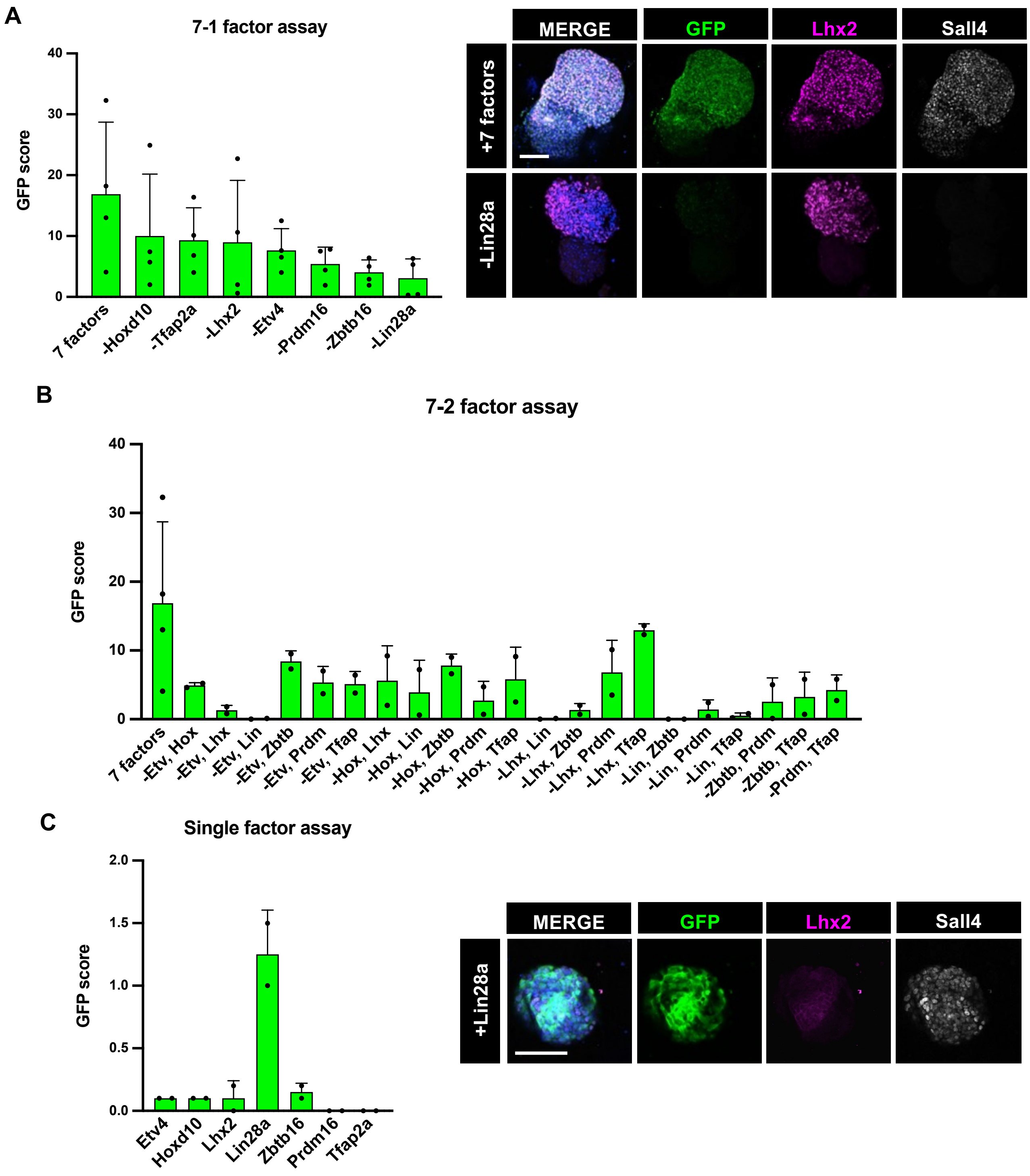


**Fig. S6 Atsuta et al.**

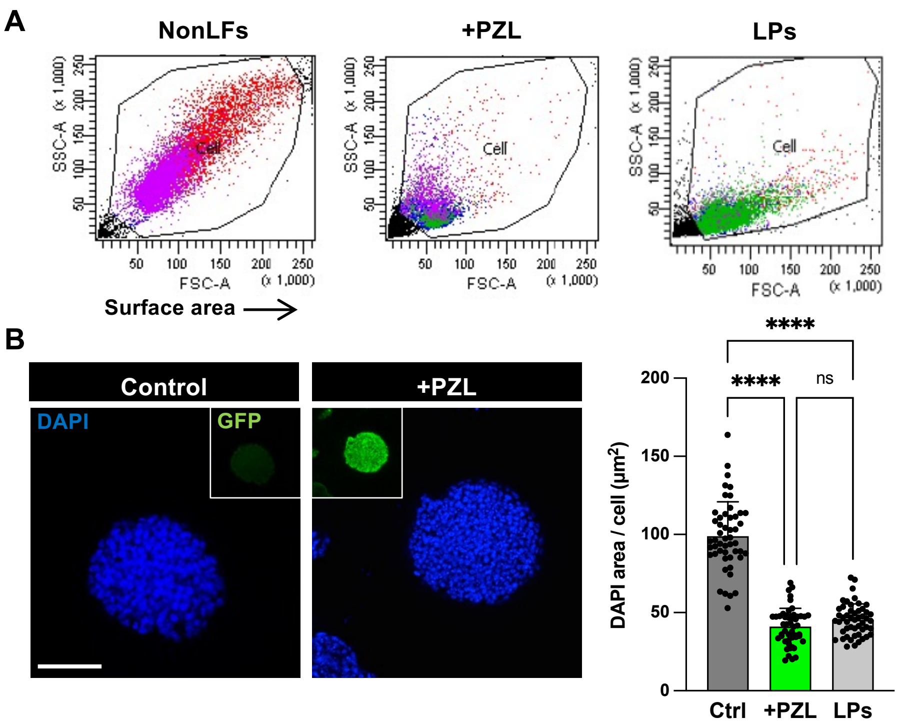


Fig. S7 Atsuta et al.

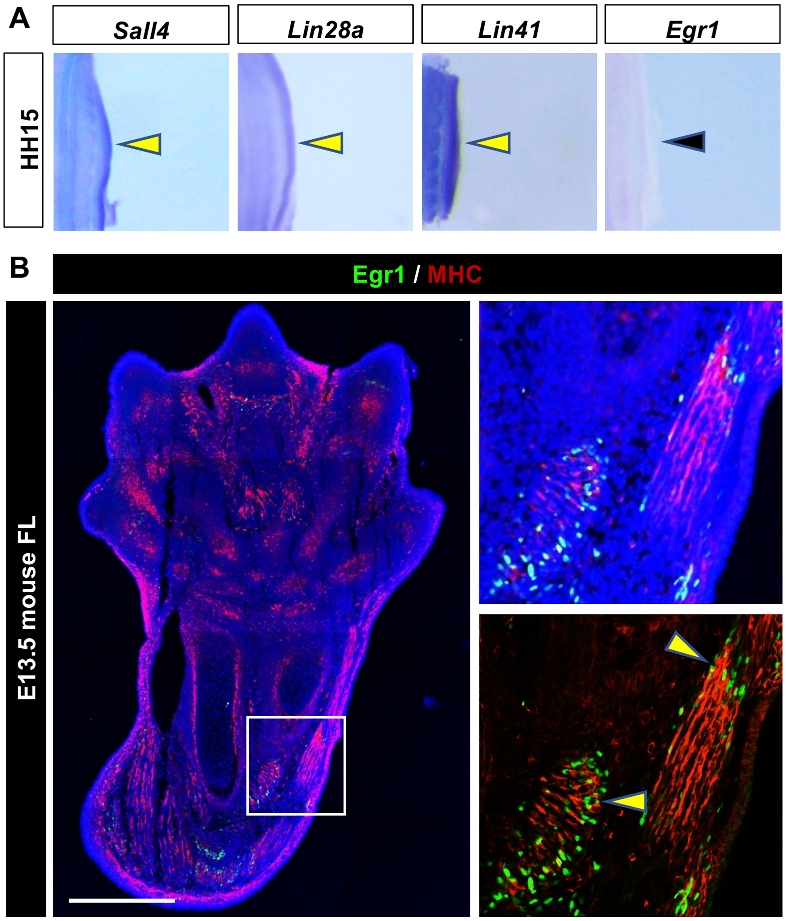


Fig. S8 Atsuta et al.

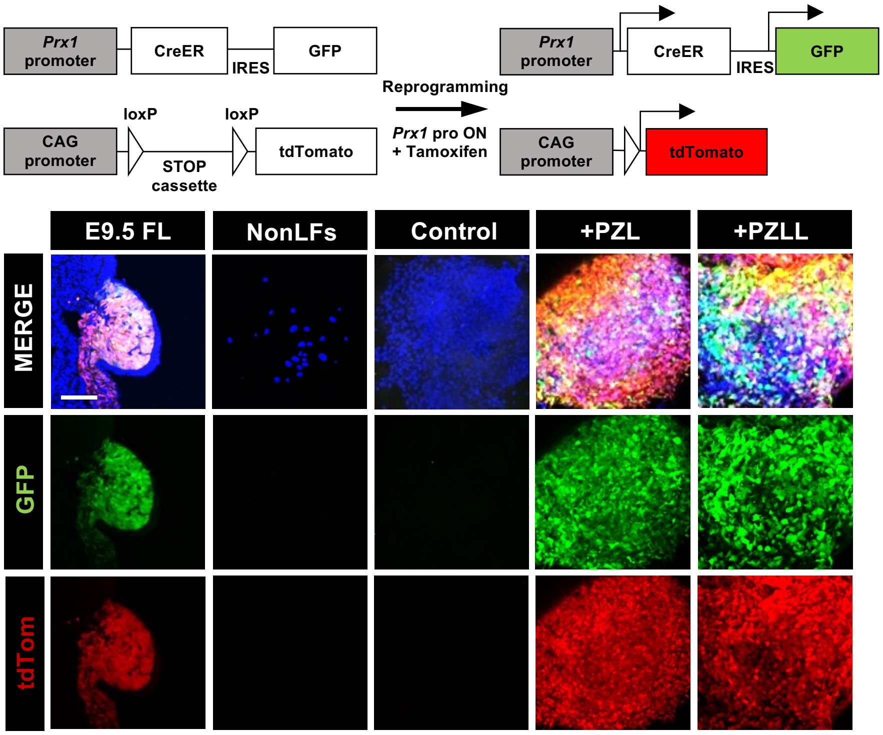


Fig. S9 Atsuta et al.

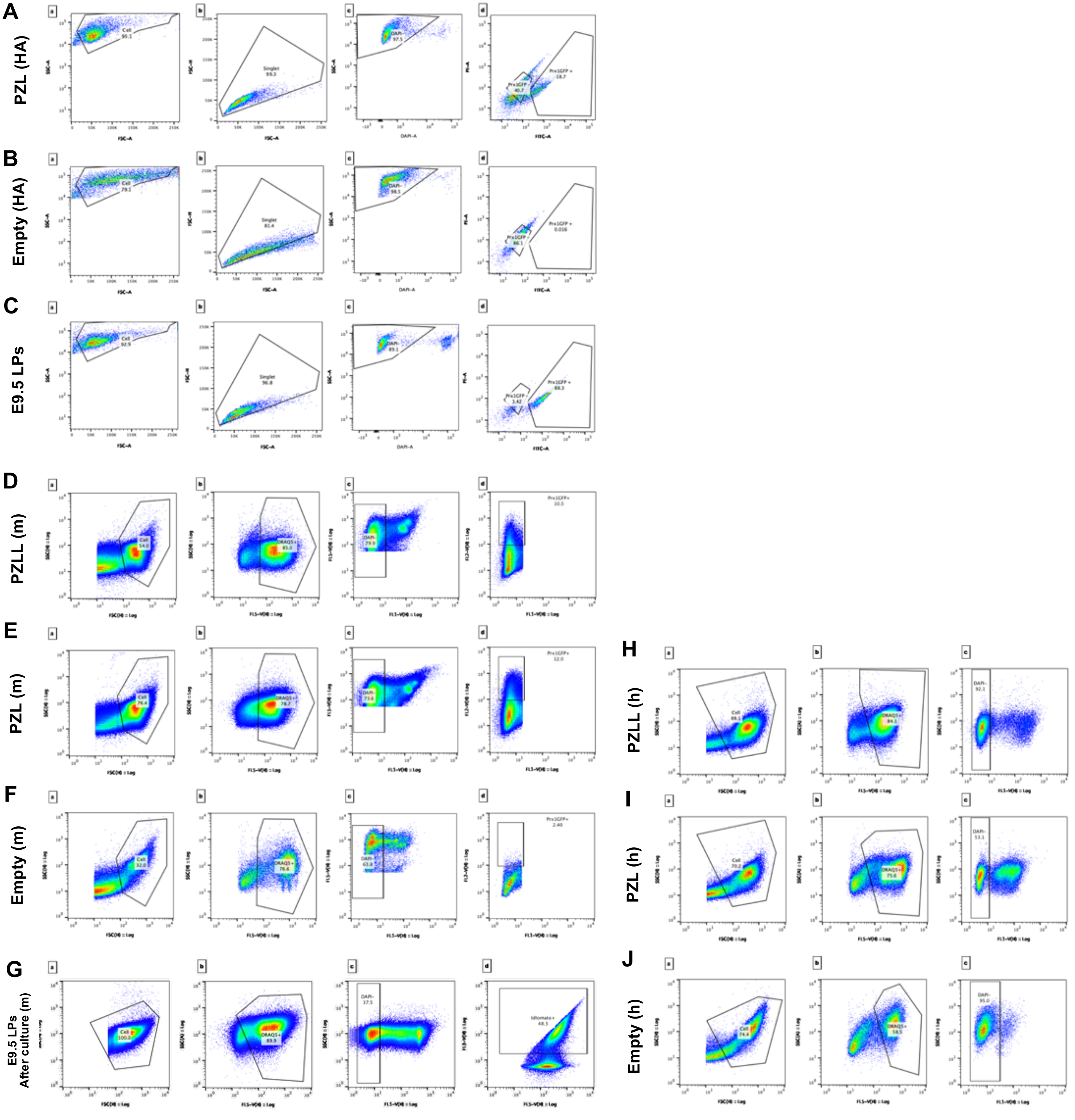


**Fig. S10 Atsuta et al.**

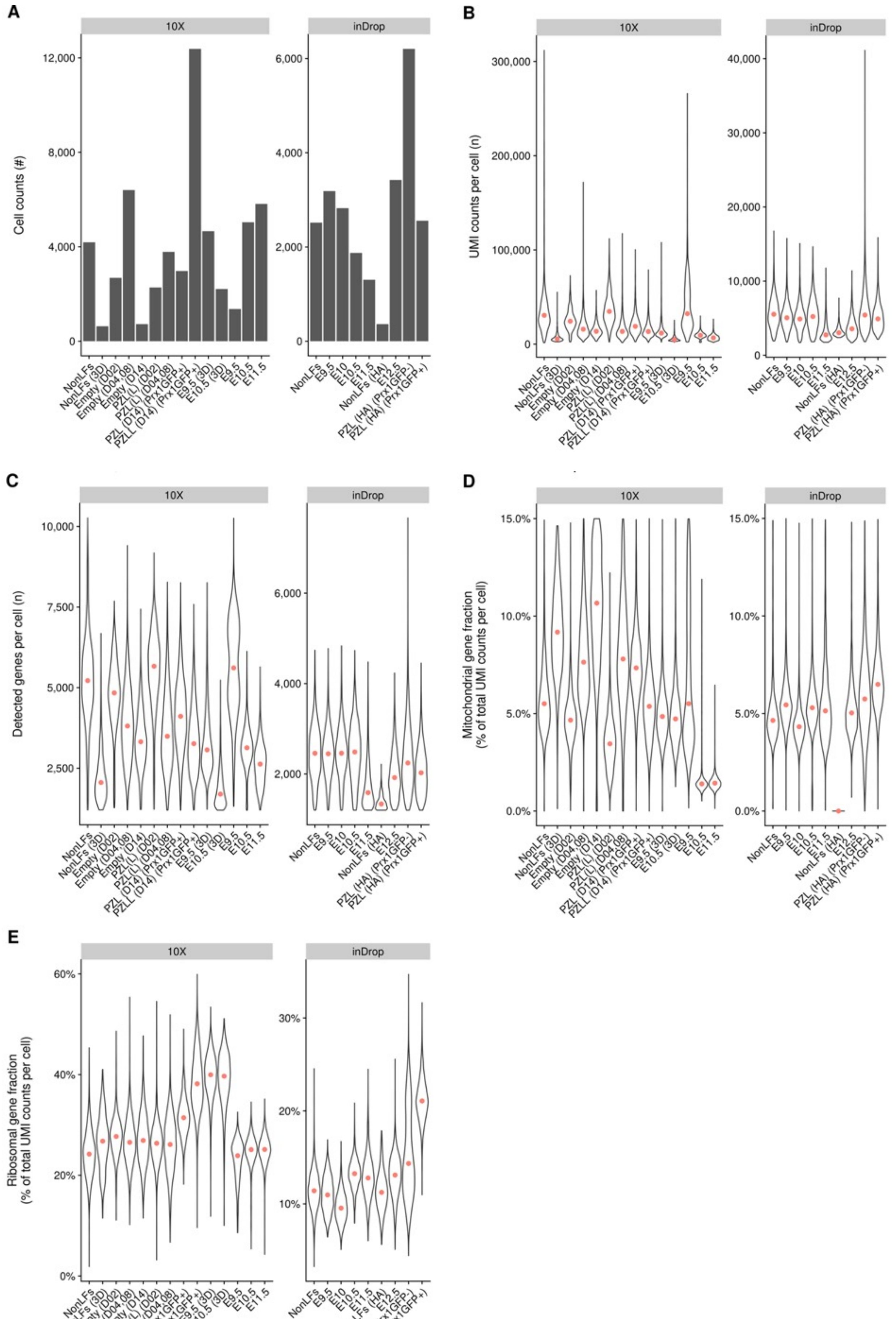


Fig. S11 Atsuta et al.

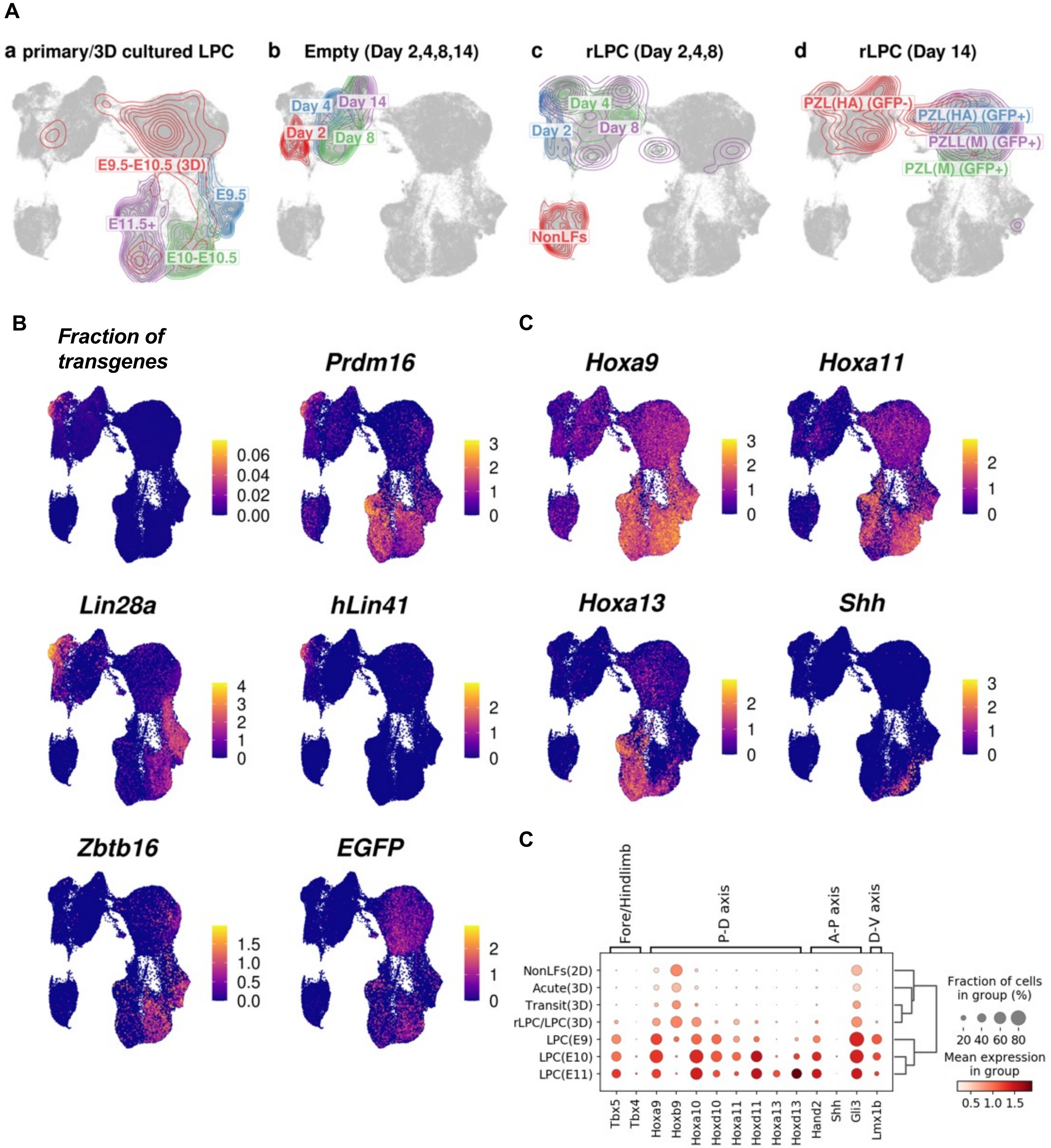


Fig. S12 Atsuta et al.

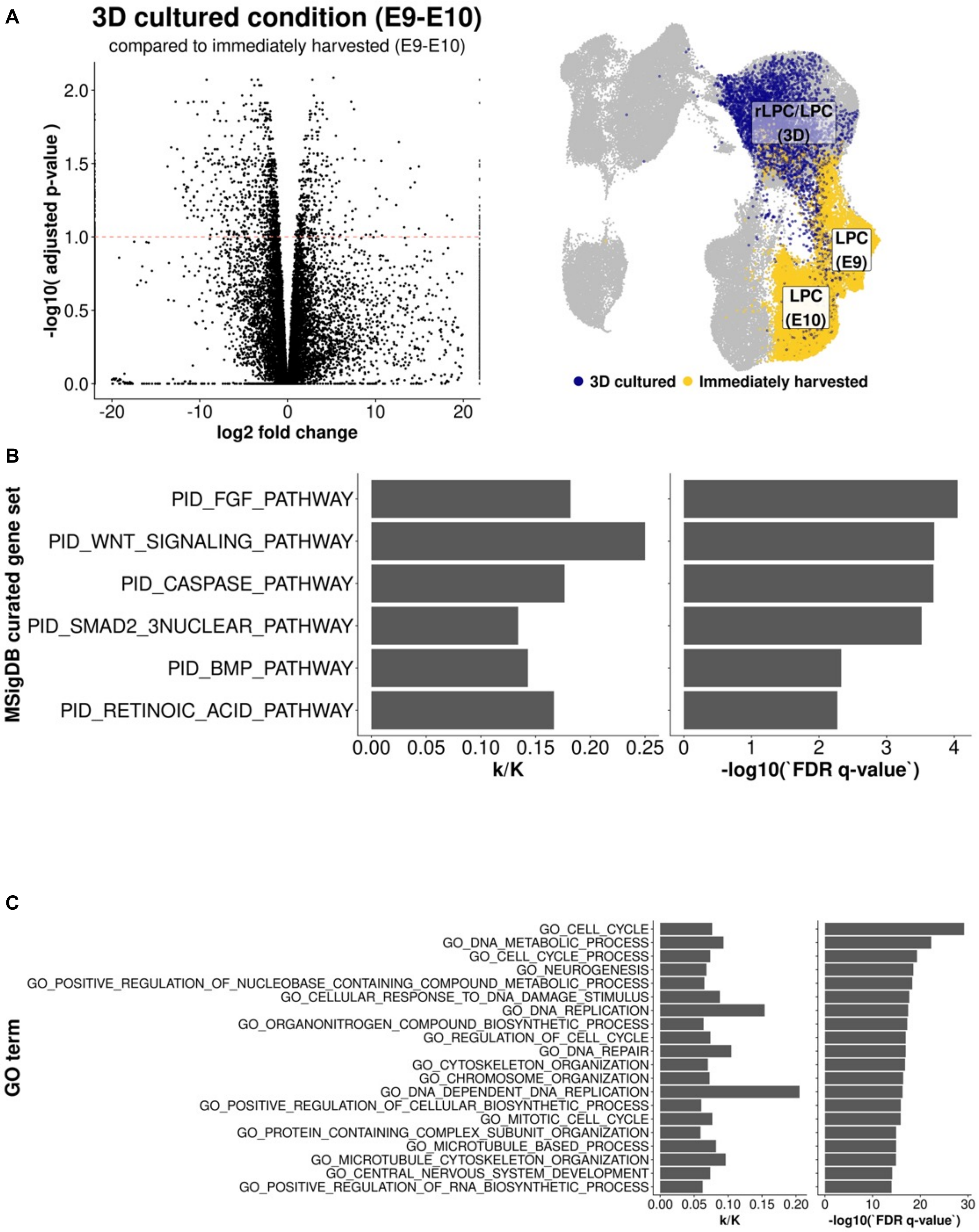


Fig. S13 Atsuta et al.

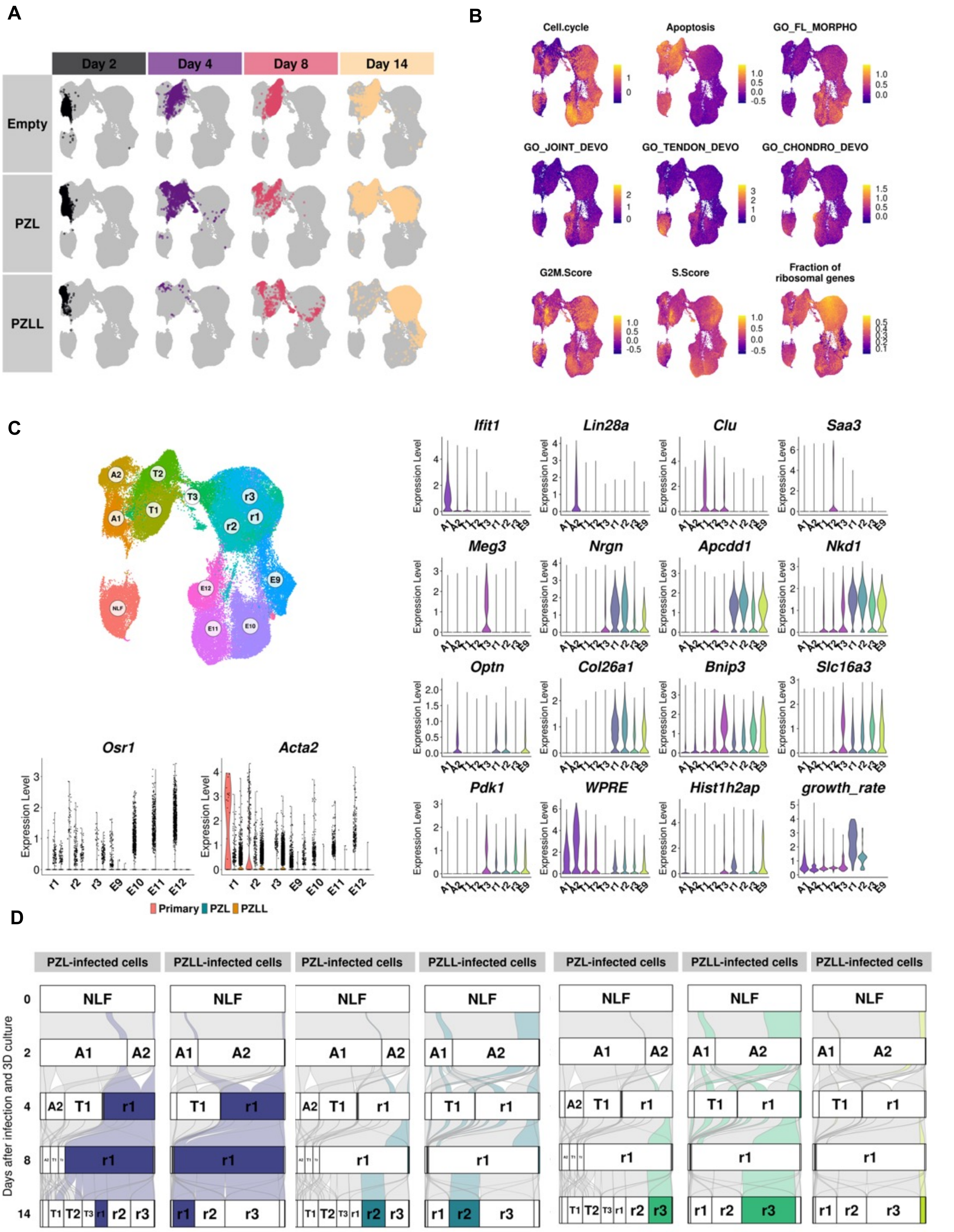


Fig. S14 Atsuta et al.

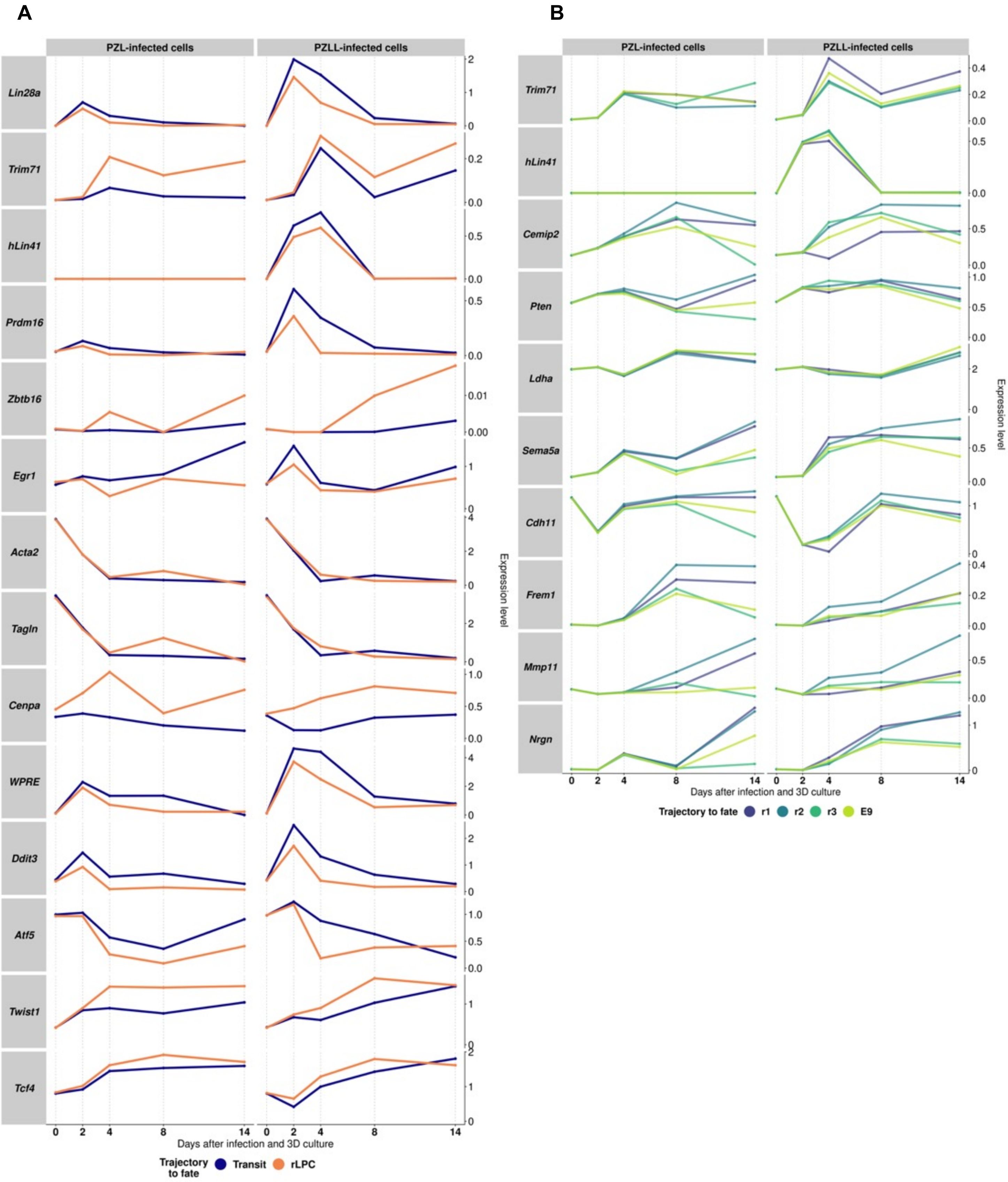


Fig. S15 Atsuta et al.

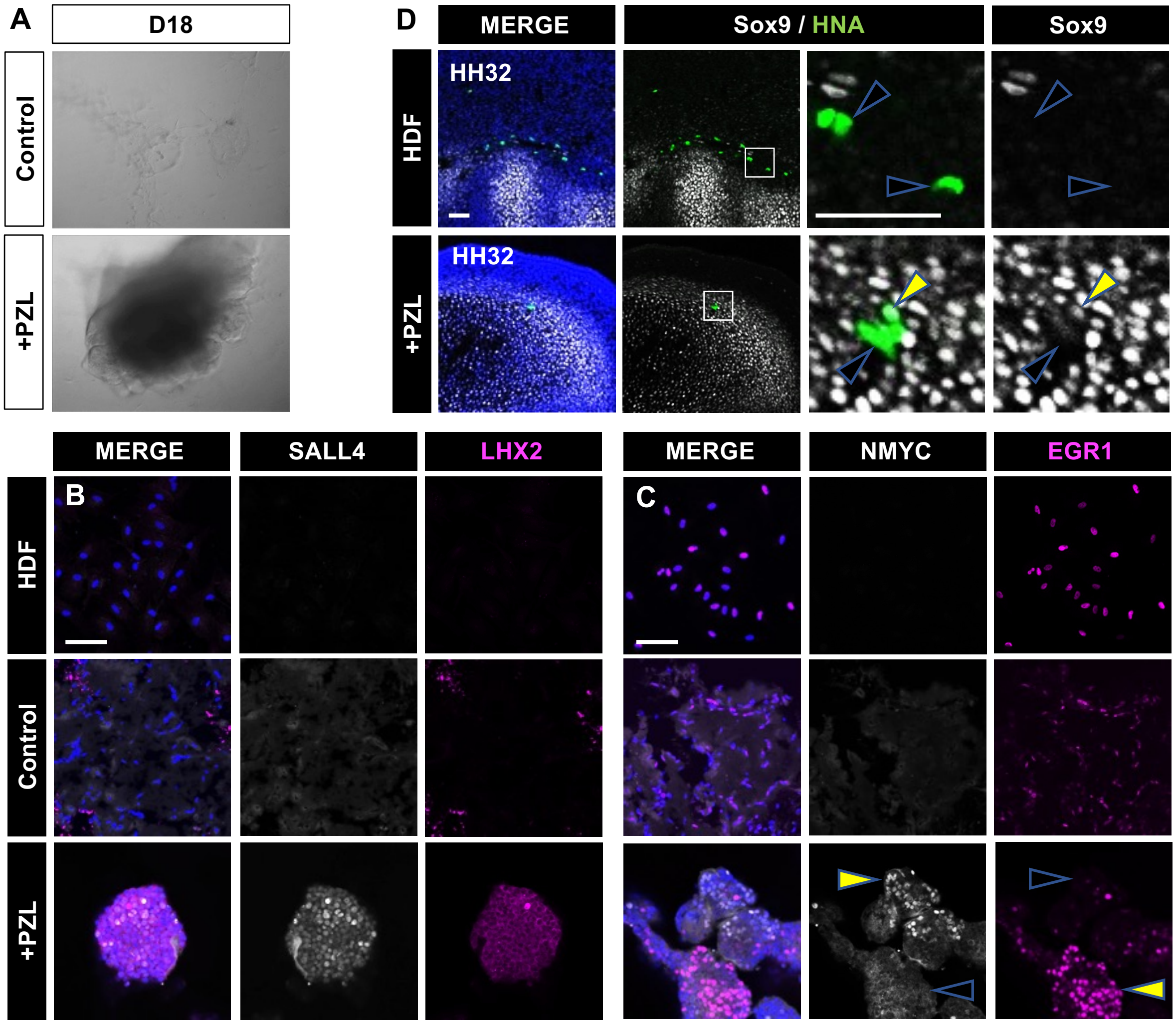


Fig. S16 Atsuta et al.

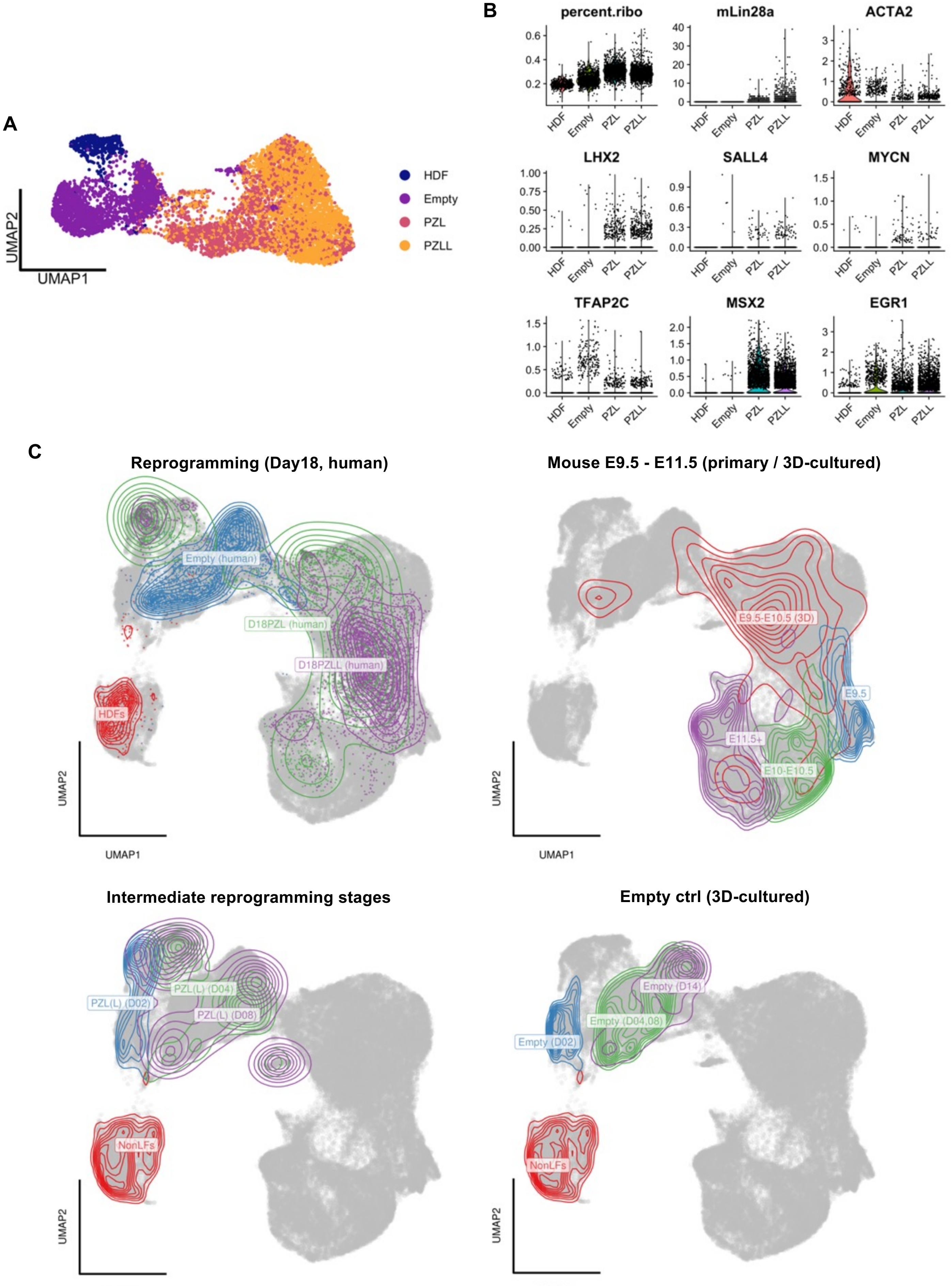


Fig. S17 Atsuta et al.
